# A neuronal substrate for translating nutrient state and resource density estimations into foraging decisions

**DOI:** 10.1101/2023.07.19.549514

**Authors:** Dennis Goldschmidt, Ibrahim Tastekin, Daniel Münch, Jin-Yong Park, Hannah Haberkern, Lúcia Serra, Célia Baltazar, Vivek Jayaraman, Gerald M. Rubin, Carlos Ribeiro

**Author notes:** authors contributed equally (Dennis Goldschmidt, Ibrahim Tastekin).

## Abstract

Foraging animals must balance the costs of exploring their surroundings with the potential benefits of finding nutritional resources. Each time an animal encounters a food source it must decide whether to initiate feeding or continue searching for potentially better options. Experimental evidence and patch foraging models predict that this decision depends on both nutritional state and the density of available resources in the environment. How the brain integrates such internal and external states to adapt the so-called exploration-exploitation trade-off remains poorly understood. We use video-based tracking to show that *Drosophila* regulates the decision to engage with food patches based on nutritional state and travel time between food patches, the latter being a measure of food patch density in the environment. To uncover the neuronal basis of this decision process, we performed a neurogenetic silencing screen of more than 400 genetic driver lines with sparse expression patterns in the fly brain. We identified a population of neurons in the central complex that acts as a key regulator of the decision to engage with a food patch. We show that manipulating the activity of these neurons alters the probability to engage, that their activity is modulated by the protein state of the animal, and that silencing these neurons perturbs the ability of the animal to adjust foraging decisions to the fly’s travel time between food patches. Taken together, our results reveal a neuronal substrate that integrates nutritional state and patch density information to control a specific foraging decision, and therefore provide an important step towards a mechanistic explanation of the cognitive computations that resolve complex cost-benefit trade-offs.

## Introduction

All animals must find and consume food to survive and thrive. The task of locating, selecting and consuming food becomes particularly challenging in complex environments with uncertain and uneven distributions of resources (Hills et al., 2015). Thus, foraging provides a rich and ethologically relevant framework to study how the brain makes decisions (Mobbs et al., 2018; Dennis et al., 2021). Importantly, foraging is guided not only by external cues, but also by the animal’s internal needs (Dethier, 1957; Dietrich et al., 2015; Corrales-Carvajal et al., 2016). One example of this phenomenon are nutrient-specific appetites where current and predicted nutritional needs are translated into changes in foraging and feeding decisions, allowing the animal to maintain nutrient homeostasis (Corrales-Carvajal et al., 2016; Simpson and Raubenheimer, 2012; Münch et al., 2020; Walker et al., 2017; Chen et al., 2015). At the circuit level, most of what we know about nutritional state-dependent modulation of behavior focuses on consummatory behavior, and therefore mainly on gustatory sensory neurons (Steck et al., 2018; Inagaki et al., 2014; Marella et al., 2012) and on how chemosensory information is used to initiate feeding (Münch et al., 2022; Snell et al., 2022; Shiu et al., 2022). How nutritional states affect higher-order processing and cognitive computations is still poorly understood.

One of the most influential formalizations of how animals might optimize foraging is patch foraging (Stephens and Krebs, 1986). Patch foraging assumes that the animal moves around in an environment with patchy distributions of food and is faced with the tasks of deciding which patches it will visit (exploration) and how much food it will ingest from each visited patch (exploitation; that is, how long it will feed on a patch). Because exploring for food patches and staying on a food patch to feed are mutually exclusive processes, animals must negotiate the so-called exploration-exploitation trade-off (Sutton and Barto, 1998) to optimize foraging (Addicott et al., 2017). This trade-off, which has been studied extensively in behavioral ecology and computational neuroscience, is thought to be central to decision-making and risk assessment in most animals (Dall et al., 2005; Addicott et al., 2017). Important algorithmic and neuronal insights into how animals solve the exploration-exploitation trade-off have come from the use of precisely controlled behavioral paradigms in rodents and primates, including humans (Kane et al., 2022, 2019; Hayden et al., 2011; Kolling et al., 2012). However, it has been difficult to mechanistically dissect these processes in naturalistic foraging settings in which the animal is allowed to freely forage in a patchy food environment. Additionally, most prior studies have focused on the decision of the animal to leave the patch (stop exploiting). Another key aspect of foraging decisions—whether to engage with the food patch and start feeding—remains largely unexplored.

Decision-making in foraging animals is affected by the cost of searching: for example, the lower the density of resources (food patches) in the environment, the higher the value of each patch (Stephens and Krebs, 1986). For this reason, animals have been shown to spend more time on patches when resource density is low than when resource density is high. One of the most successful optimality models explaining this phenomenon is known as the Marginal Value Theorem (MVT) (Charnov, 1976). The MVT postulates that an animal adjusts how long it feeds on a patch depending on the time that it takes to travel between patches, implying that an animal adjusts its foraging decisions based on its internal estimate of resource density. Indeed, many studies have shown this qualitative relationship holds in animals ranging from vertebrates—like mammals (Cassini et al., 1990) and birds (Cowie, 1977; Kacelnik, 1984)—to insects (Wajnberg et al., 2000; Bonser et al., 1998; Kay, 2002; Parker, 1992). While the original assumption of the MVT is that the costs of searching mainly relate to the energy cost of moving, other factors such as opportunity cost and the risk of losing track of a food patch with high nutrient content might also play a role.

Foraging decisions require the integration of past experience, internal states, and information from multiple sensory modalities to infer the statistics of the environment (Stephens et al., 2007). In insects, the central complex (CX) is a higher-order brain neuropil that is suitable for such complex decision-making (Strauss, 2002; Pfeiffer and Homberg, 2014; Turner-Evans and Jayaraman, 2016; Heinze, 2017; Honkanen et al., 2019). Recent studies have identified and characterized all fly CX neuronal cell types as well as their connectivity (Wolff et al., 2015; Wolff and Rubin, 2018; Hulse et al., 2021). Therefore, the CX has become a tractable neuropil for studying the integration of sensory and internal state signals with the aim of understanding how adaptive behavioral policies are generated (Seelig and Jayaraman, 2015; Kim et al., 2017; Green et al., 2017; Giraldo et al., 2018; Okubo et al., 2020; Dan et al., 2021; Matheson et al., 2022). While significant progress has been made in understanding navigational and vector computations in the fly (Seelig and Jayaraman, 2015; Kim et al., 2017; Green et al., 2017; Stone et al., 2017; Hulse et al., 2021; Lyu et al., 2022; Lu et al., 2022; Fisher, 2022) much less is known about how complex non-navigational decisions are controlled by the CX. Furthermore, the CX also integrates metabolic states to regulate sleep (Donlea et al., 2014; Liu et al., 2016; Donlea et al., 2018) and feeding (Sareen et al., 2021; Park et al., 2016; Musso et al., 2021). The precise cellular identity of the neurons supporting these behavioral adaptations and which aspects of the overall behavioral strategies are altered to ensure homeostasis remains to be identified.

Here, we performed a large-scale neuronal silencing screen to identify neuronal subsets involved in foraging decisions. We first built an experimental setup for highthroughput video-based tracking of individual flies foraging in an arena with yeast and sucrose patches. This allowed us to study the effect of silencing neurons labeled in more than 400 GAL4 driver lines, revealing a neuronal population projecting to the dorsal fan-shaped body (dFB), a layered structure within the CX, as being important for the transition from exploration to exploitation. Silencing these neurons leads to an increase in the probability to engage with food patches, while other foraging parameters and overall food intake remain unaffected. Likewise, acute optogenetic activation of these neurons as the animal approached a food patch led to a reduction in the probability to engage with food patches and initiate feeding. We then investigated which neuronal subsets of this population were responsible for the phenotype and showed that a single pair of dFB neurons (FB6A.1) is required to suppress the initiation of feeding upon encountering food. Strikingly, whole-cell patch-clamp recordings showed that the firing rate in the FB6A.1 neurons is modulated by the protein state of the fly, such that protein deprivation decreases the firing rate of these neurons. Finally, we showed that wild-type flies modulate their probability to engage with food patches based on their inferred density. Importantly, silencing dFB neurons abolishes the relationship of travel time between food patches with the probability to engage upon food encounter—a computation which has been proposed to underlie the adaptation of foraging decisions to resource distributions in classical optimal foraging models (Charnov, 1976). Our work identifies a specific higher-order neuronal substrate that plays a role in integrating both internal nutrient needs and external nutrient availability distributions to regulate a behavioral policy underlying transitions from exploring the environment to exploiting an encountered food patch.

## Results

### High-throughput image-based behavioral tracking for dissecting foraging decisions

Naturalistic behaviors performed by freely moving animals are notoriously challenging to capture and describe quantitatively (Dennis et al., 2021; Kennedy, 2022). Recent advances in software and hardware technologies allow for high-resolution, and high-throughput quantitative analysis of the behavior of freely moving animals (Branson et al., 2009; Bath et al., 2014; Alisch et al., 2018; Barlow et al., 2022; Mathis et al., 2018). Inspired by an existing setup (Corrales-Carvajal et al., 2016) developed to quantify foraging decisions performed by *Drosophila melanogaster*, we built an automated image-based tracking setup that captures trajectories of an individual fly in a foraging arena with 6 radially distributed and interspersed patches of yeast (amino acid source) and sucrose (carbohydrate source) (Fig. 1a, Suppl. Video 1, Suppl. Fig. 1a, also see Methods). We optimized the setup for scalability, robustness, and modularity in order to make it suitable for highthroughput neurogenetic behavioral screens (Suppl. Fig. 1b, also see Methods for details).

**Figure 1.**
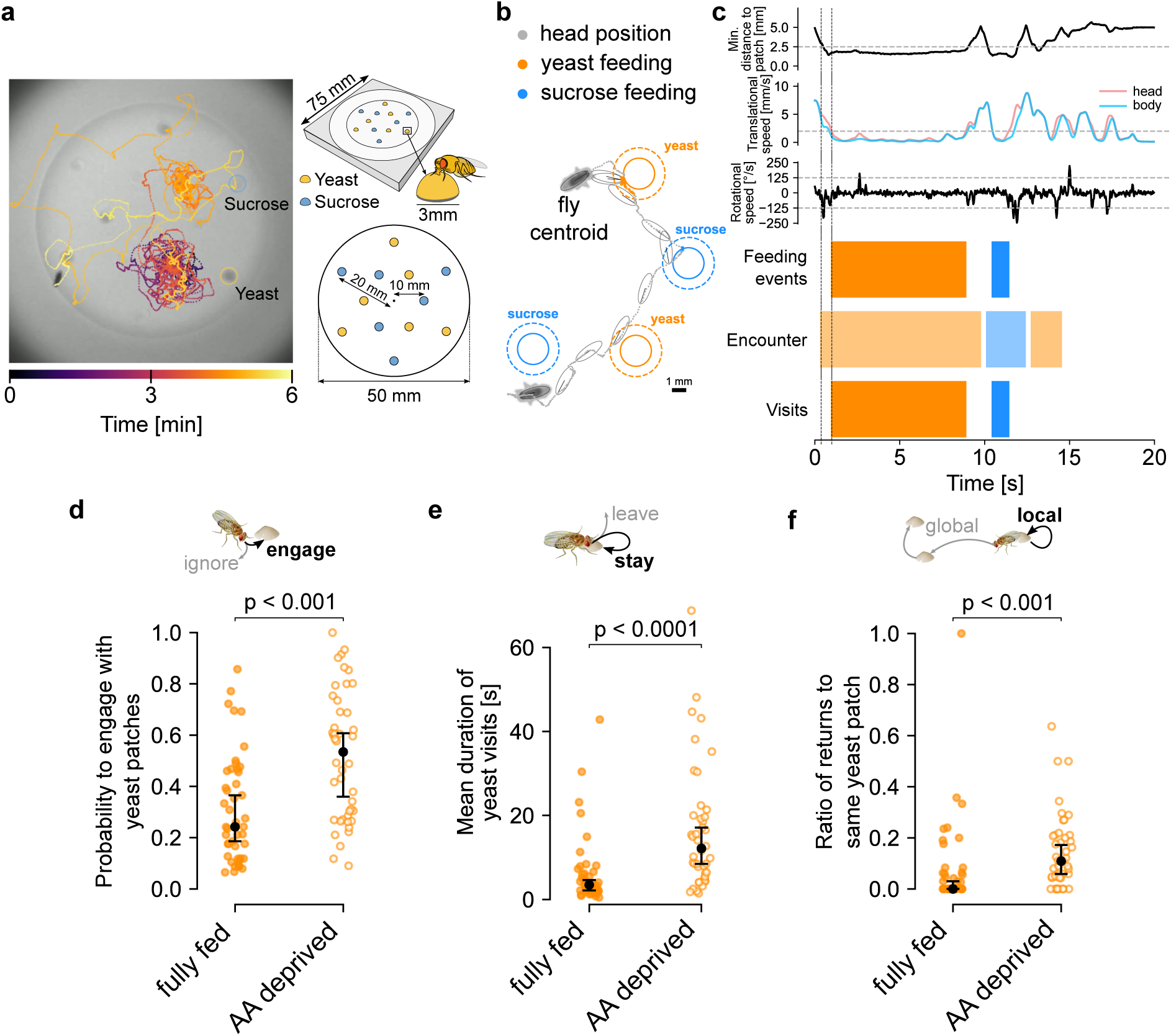
**High-throughput image-based behavioral tracking for dissecting foraging decisions. a**, Left: trajectory of a fly foraging in an arena containing sucrose and yeast patches. Right: schematic of the patch foraging arena used in this study. **b**, Automated analysis pipeline extracted the trajectories of the fly’s head and body centroid over time as it forages and feeds on yeast and sucrose patches. **c**, Time-series analysis of the distance from the fly’s head to the closest patches, translational and rotational speed by means of thresholding is used (top) for automated detection of patch feeding events, encounters and visits (bottom) using the thresholds defined by the dotted lines in the time-series plots (top). **d**, Probability to engage with yeast patches for fully fed and AA-deprived flies. **e**, Mean duration of yeast visits in seconds (s) for fully fed and AA-deprived flies. **f**, Ratio of returns to the same yeast patch for fully fed and AA-deprived flies. **d-f**, Data points are individual flies, black dots and bars indicate median, and extrema within 1.5 of the inter-quartile range (IQR), respectively. P-values are obtained by performing Wilcoxon rank-sum tests, *n =*44-45.

From the tracked trajectories, we extracted the fly’s distance to the closest food patch and the time series for various kinematic parameters, such as the fly’s rotational speed and the translational speeds of its head and body (Fig. 1b and c, see also Methods for details) (Corrales-Carvajal et al., 2016). Previous behavioral studies have shown that slow micromovements of the head near a food patch can be used as a proxy for feeding (Corrales-Carvajal et al., 2016; French et al., 2021), a measure we employed as well (Fig. 1c). We additionally defined multiple feeding events on the same patch as a “visit” to that food patch. Finally, when a fly’s head was close enough to a food patch to be in contact with it (distance < 3 mm), we defined these as “encounters”. Importantly, although both encounters and visits might include bouts in which flies touch food patches, only visits feature feeding.

### Flies adapt different foraging decisions to their nutritional state

Foraging can be broadly divided into exploration and exploitation phases (Stephens and Krebs, 1986). While exploration is key for the animal to identify the distribution and content of different food patches, it prevents the animal from staying on a food patch to ingest significant amounts of food (exploitation). The ability of animals to balance this exploration-exploitation trade-off is therefore crucial to matching foraging outcomes with physiological needs to efficiently achieve nutrient homeostasis (Münch et al., 2020). In simple terms, upon encountering a food patch, the animal balances this trade-off by deciding to either stop and engage in feeding or continue exploring. The decision to engage with a food patch or not is therefore central to negotiating the exploration-exploitation trade-off. To capture this decision, we calculated the ratio of the total number of visits to the total number of encounters. We then used this effective probability to engage with a food patch as our metric for how flies balance the exploration-exploitation trade-off (Corrales-Carvajal et al., 2016).

Within foraging theory, multiple quantitative frameworks, of which the budget theory is the most famous, predict that foraging behavior should be modulated by the nutritional state of the animal (Stephens, 1981; Caraco et al., 1980; Kacelnik and El Mouden, 2013). Consistent with the results of an earlier study in which we showed that flies that are deprived of amino acids (AA) specifically alter different foraging parameters to restore nutrient homeostasis (Corrales-Carvajal et al., 2016). We observed that AA-deprived animals in our setup specifically increase their feeding on yeast, a source of AAs (Suppl. Fig. 2a and b). This is the probability to engage with yeast patches (Fig. 1d) is also increased by AA deprivation, and AA-deprived flies are significantly less likely to leave yeast patches (as quantified by an increase in their mean duration of yeast visits) (Fig. 1e). These effects are nutrient-specific as the behavior of AA-deprived flies towards sucrose patches remained unaltered (Suppl. Fig. 2c and 3) and are likely to strongly contribute to the specific increase in yeast feeding. Conversely, carbohydrate-deprived flies specifically bias their exploitative behavior towards sucrose patches and not yeast (Suppl. Fig. 4). We also quantified how flies explore the arena. As a measure of how local the exploratory pattern of a given fly is, we analyzed how flies transition from one yeast patch to the next by comparing the number of returns to the same yeast patch to the total number of transitions between yeast patches. Similar to previous results, we observed that AA deprivation increases the ratio of returns to the same yeast patch (Fig. 1f); the AA-deprived fly exhibits a more local exploration pattern, which can be interpreted as reduced risk-taking by remaining close to a known food source, while still searching for better food sources (Suppl. Video 1). This nutritional-state modulation is also evident in the reduced area explored by AA-deprived flies within defined time bins (Suppl. Fig. 5). These results demonstrate that we can quantify distinct decision-making processes that regulate foraging, as well as how they are modulated by the animal’s nutritional state, using the highthroughput screening methods we established.

### A neurogenetic silencing screen identifies neuronal subsets required for the decision of transitioning from exploration to exploitation

To identify neurons that regulate foraging decisions, we performed a large-scale neuronal silencing screen, in which we inactivated neuronal subsets labeled by different GAL4 driver lines in the central brain and measured the effects of silencing on foraging parameters defined in the previous section. We focused on the decision to engage and start feeding at a food patch (measured as the probability to engage with a yeast patch) due to its relevance for how the animal balances the exploration-exploitation trade-off. We selected 434 GAL4 driver lines from the Janelia FlyLight Generation 1 GAL4 collection (Jenett et al., 2012) with a focus on lines with a sparse expression pattern, which targeted a wide repertoire of brain regions (Suppl. Fig. 6). We inactivated neurons labeled in the GAL4 lines by inducing the expression of the inward-rectifier potassium channel Kir2.1 in a temperature-dependent manner in adults for one to two days prior to screening in order to minimize any developmental effects that might be caused by the constitutive expression of Kir2.1 (Baines et al., 2001). Protein-deprived flies with silenced neurons were recorded for one hour while foraging in the arena containing yeast and sucrose patches (Fig. 1a). We tested each line and its corresponding genetic controls within the same batch and calculated the probability to engage with a yeast patch. The resulting distribution for this behavioral parameter was centered around the median of all control flies tested and spanned a wide phenotypic range of values with an enrichment for lines showing an increase in the probability to engage (Fig. 2a). We then performed a bootstrapping analysis for phenotypes deviating significantly from the whole-screen population distribution leading to the identification of 49 lines (Fig. 2b and c, see Methods for details). 32 of these showed a significant difference to the control lines tested within the same batch (Fig. 2c). Based on multiple rounds of retests, 3 lines were selected for further characterization as they showed a reproducible increase in the probability to engage with a yeast patch (Fig. 2d and Suppl. Fig. 7). Other foraging parameters, such as the mean duration of yeast visits, that is, the decision to leave a patch (Fig. 2e and Suppl. Fig. 8a and d), and the ratio of returns to the same yeast patch, that is, the spatial organization of exploration (Fig. 2f and Suppl. Fig. 8b and e), were not consistently affected upon silencing the neurons labeled by these GAL4 driver lines. Thus, the silencing phenotypes are specific to the transition from exploration to exploitation, but not other general aspects of the exploration or exploitation phases. Furthermore, the total duration of micromovements (a proxy for total food intake) (Fig. 2g, Suppl. Fig. 8c and f) was not consistently affected by the silencing, strongly suggesting that the labeled neurons are not involved in setting the overall nutritional state of the fly. The absence of a change in overall food intake was further corroborated by using the flyPAD assay (Itskov et al., 2014), which directly measures the total amount of feeding (Suppl. Fig. 9). Given the specificity of the phenotype these data suggest that different foraging decisions are controlled independently by distinct neuronal populations. Taken together, these observations indicate that these three GAL4 lines label neurons important for mediating the decision to engage with a yeast patch as part of the process of balancing the explorationexploitation trade-off.

**Figure 2.**
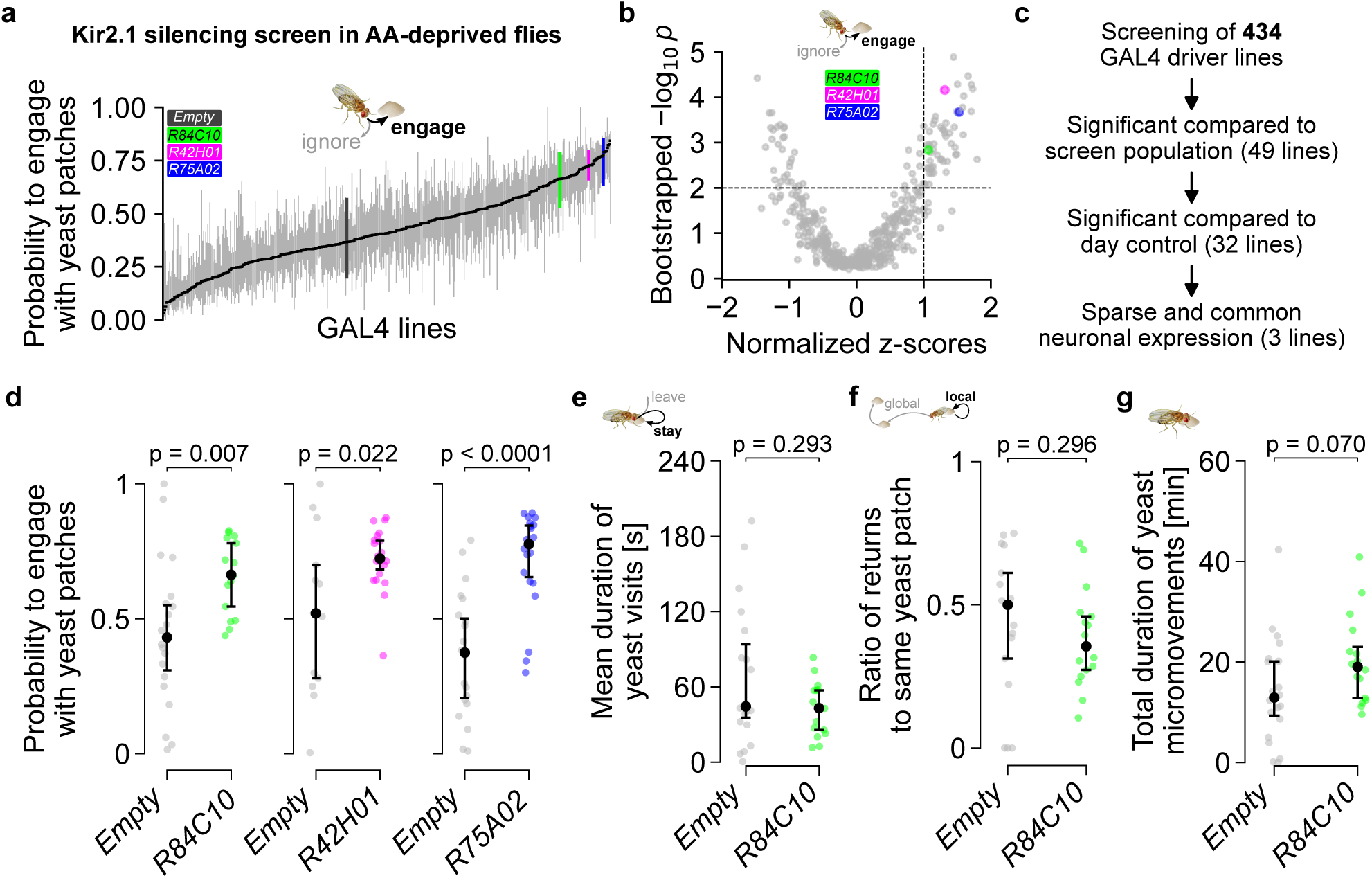
**A neurogenetic silencing screen identifies GAL4 lines producing specific phenotypes in the decision to transition from exploration to exploitation. a**, Distribution of the probability to engage with yeast patches upon silencing of neurons labeled in 434 different GAL4 driver lines. The distribution is sorted by median values indicated by the horizontal black line. Gray outlines are IQR of the first and third quartile for each line. Control flies (*Empty-GAL4* black) and specific GAL4 lines (*R84C10*, *R42H01*, *R75A02*) are marked with the corresponding colors. **b**, Volcano plot of normalized z-scores vs. bootstrapped p-values for the probability to engage with yeast patches for each tested GAL4 line. Thresholds used for systematic identification of hit driver lines are indicated by dashed lines and the three GAL4 lines chosen for further analysis are labeled with the corresponding colors. **c**, Diagram of the decision process used to identify positive GAL4 driver lines. **d**, Silencing phenotype in the probability to engage with yeast patches observed in the experimental GAL4 lines when compared to the *Empty-GAL4* control flies run in parallel in the same batch. **e-g**, Silencing phenotypes for other foraging parameters observed in *R84C10-GAL4* and corresponding control flies: (**e**) mean duration of yeast visits (**f**) ratio of returns to the same yeast patch and (**g**) total duration of yeast micromovements. Dot and error bars in (**d-g**) indicate medians and IQR, data points correspond to individual flies. P-values are obtained by performing Wilcoxon rank-sum tests, *n =*14-20.

### Dorsal fan-shaped body neurons specifically control the decision to engage with food patches during foraging

To identify the brain regions involved in controlling the probability to engage, we compared the labeling pattern of the three identified GAL4 lines—*R84C10*, *R42H01*, and *R75A02*—in the brain and the VNC (Fig. 3a and Suppl. Fig. 10). *R84C10* showed the sparsest expression with all three lines labeling a set of neurons that seemed to overlap in the dorsal fan-shaped body (dFB) of the CX. To better compare their expression, we overlaid the registered expression patterns of these driver lines on a template brain (JRC2018Unisex; Court et al., 2023; Bogovic et al., 2020) (Fig. 3b), which revealed that they indeed had overlapping projections to the FB layer 6 (FB6) and the Superior Neuropils (SNP), with cell bodies located in the lateral posterior part of the protocerebrum.

**Figure 3.**
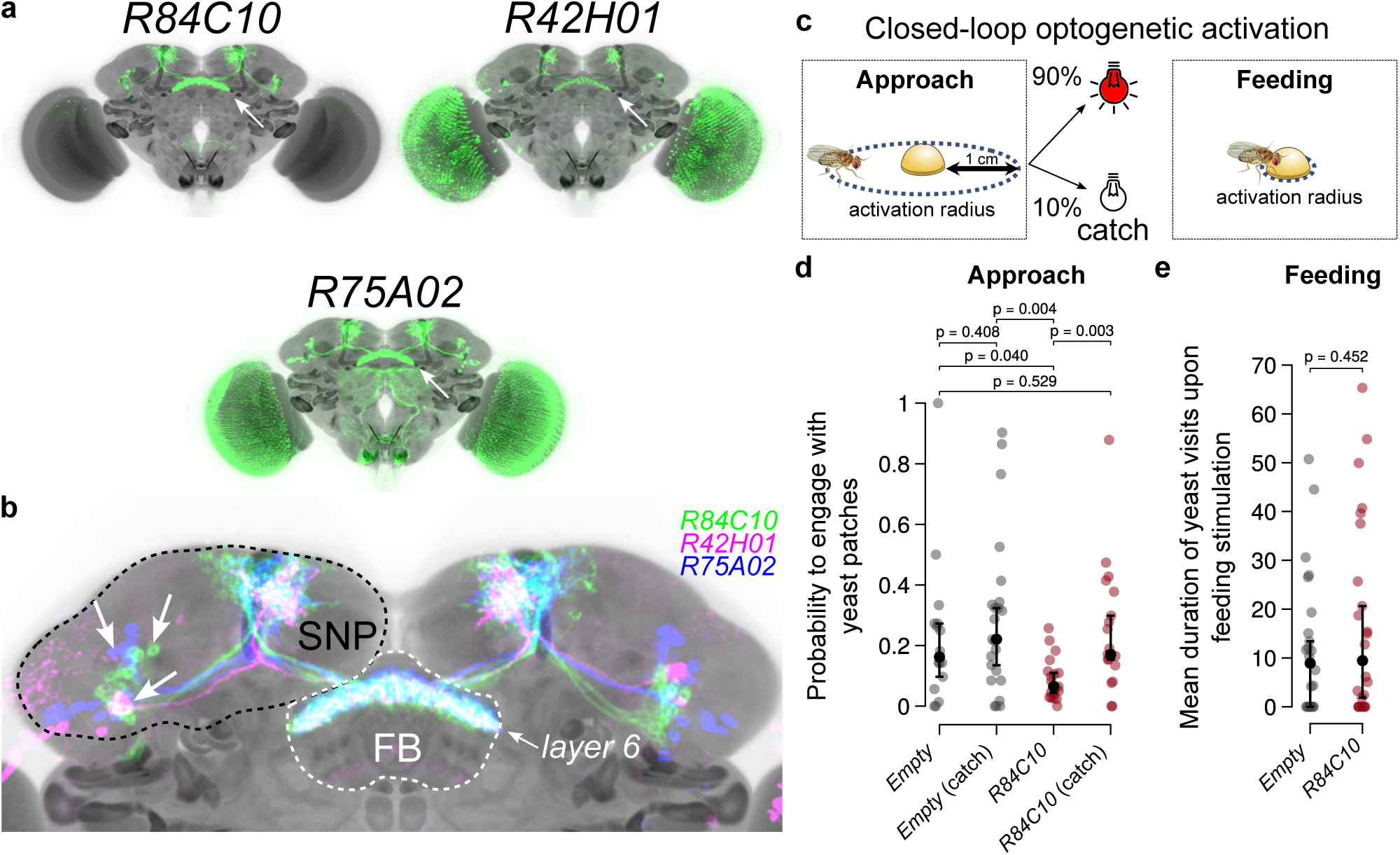
**The identified GAL4 lines label the same dorsal fan shaped body layer and their activation is sufficient to decrease the probability to engage with yeast patches. a**, Aligned maximum intensity projection confocal stacks of *R84C10-*, *R42H01-*, and *R75A02-GAL4* driving mCD8::GFP expression reveals neurons projecting to the dorsal fan-shaped body (dFB, white arrows). **b**, Overlapping the expression patterns of *R84C10-*, *R42H01-*, and *R75A02-GAL4* reveals common labeling of neurons projecting to layer 6 of the FB neurons with soma positioned in the lateral posterior brain and projections to the Superior Neuropils (SNP). **c**, Experimental protocol for closed-loop optogenetic activation in AA-deprived flies while the fly is approaching (left) or feeding on (right) a yeast patch. 10% of approaches do not lead to the activation of the LED (catch trials). **d**, Effect of genotype and optogenetic protocol on the probability to engage with yeast patches for the approach experiments. e, Effect of optogenetic stimulation while feeding on the mean duration of yeast visits. Dot and error bars in (**d-e**) indicate medians and IQR, data points correspond to individual flies. P-values are obtained by performing Wilcoxon rank-sum tests, *n =*19-23.

Previous studies have implicated dFB neurons in regulating sleep. However, silencing dFB lines that target these distinct neural populations (*R23E10-*, *104y-*, and *c205-GAL4*), all of which have been shown to modulate sleep in flies (Donlea et al., 2014, 2018), did not lead to a phenotype in the probability to engage with yeast patches (Suppl. Fig. 11). This suggests that changing or perturbing sleep through dFB neuronal silencing is not sufficient to induce the foraging phenotypes we observed. Taken together, these results demonstrate that dFB neurons labeled by *R84C10*, *R42H01* and *R75A02* modulate the probability that the fly transitions from exploration to exploitation without affecting other foraging decisions.

### Activating dFB neurons specifically reduces the probability of the fly to transition to exploitation

Our silencing results suggest that activity in dFB neurons suppresses the transition from exploration to exploitation. Therefore, we decided to test whether dFB neurons are sufficient to decrease the probability to engage with a food patch. To do so, we performed acute optogenetic activation of these neurons in AA-deprived flies using red-shifted channelrhodopsin (Klapoetke et al., 2014). Specifically, we used a closed-loop stimulation paradigm, in which the LED is turned on as the fly either (1) approaches the food patch, (2) or has engaged in feeding (see Fig. 3c, Methods for details). Moreover, 10% of approaches did not trigger the activation of the LED (catch trials). Activating dFB neurons while the fly approached a yeast patch led to a decreased probability to engage with a yeast patch when compared to the genetic control and catch trials (Fig. 3d). This is supported by an inspection of fly trajectories, which revealed that upon stimulation, experimental flies did not reach the food patch, an effect that is not observed in control flies (Suppl. Fig. 12, Suppl. Video 2). In contrast, activation of the labeled neurons when the fly was located on a food patch and had already initiated feeding did not lead to significant differences in the duration of yeast patch visits (Fig. 3e). This suggests that the physiological role of dFB neurons is strongly dependent on the behavioral state of the animal. Moreover, consistent with our silencing experiments, the labeled neurons appear to affect only the probability that the fly transitions from exploration to exploitation, and not the duration of time spent exploiting the yeast patch. The finding that neuronal silencing and activation phenotypes produce opposite effects further supports our proposal that dFB neurons have a specific role in modulating the probability to transition from exploration to exploitation.

### A single FB6 neuron pair is necessary for adapting the probability to engage with food

Our data raise the exciting possibility that the three identified lines label a common set of neurons projecting to the FB layer 6, which specifically controls the probability of the fly to engage with a yeast patch. However, because these genetic driver lines also exhibit expression in multiple dFB cell types as well as in cells in other brain areas, such as the SEZ and VNC (Fig. 3a and Suppl. Fig. 10), further experiments were required to unambiguously assign the observed phenotypes to specific FB6 neurons.

We employed a split-GAL4 strategy (Luan et al., 2006; Pfeiffer et al., 2010; Dionne et al., 2018) to genetically access a sparser subset of the dFB neurons labeled by the GAL4 lines identified in our silencing screen. For this, we created intersectional Split-GAL4 combinations based on two of our three identified dFB GAL4 lines. We found that intersection of the *R42H01* and *R84C10* enhancers resulted in a strong expression in a single pair of neurons projecting to the FB layer 6 (Fig. 4a and b). Silencing these neurons led to a significant increase in the probability to engage with yeast patches, as we had observed upon silencing all neurons labeled in either *R42H01-* or *R84C10-GAL4* lines (Fig. 4c and e, Fig. 2d). The magnitude of the phenotype was slightly smaller, likely due to the sparser and weaker expression of GAL4 in the splitGAL4 lines. Furthermore, like the broader lines, silencing resulted in a specific phenotype in the decision to engage. Other foraging decisions, such as deciding to leave a food patch, were unaffected (Fig. 4d and f, Suppl. Fig. 13).

**Figure 4.**
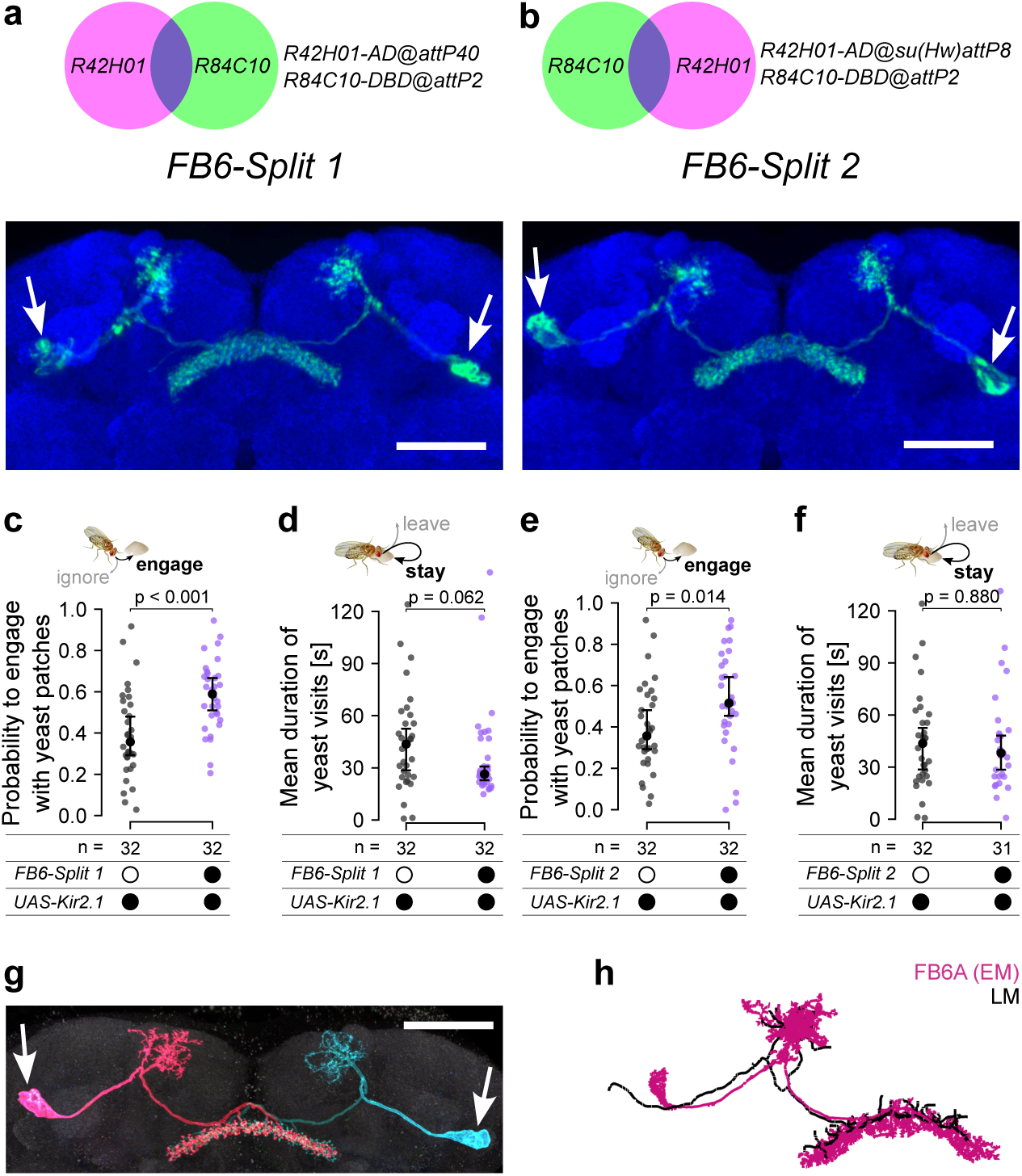
**A single pair of FB6A.1 neurons is required to modulate the probability to engage with yeast patches. a-b**, Genotype combinations of the lines used to label one FB6A.1 neuron (top) and maximum intensity projections of confocal stacks for *FB6 Split-GAL4* line 1 and 2 (bottom). Scale bar = 50 µm. **c** and **e**, Effect of silencing FB6A.1 or control genotype on the probability to engage with yeast patches. **d** and **f**, Effect of silencing FB6A.1 on the probability to engage with yeast patches. Dot and error bars in (**c-f**) indicate medians and IQR, data points correspond to individual flies. P-values are obtained by performing Wilcoxon rank-sum tests, *n =* 31-32. **g**, Maximum intensity projection of a confocal stack for FB6A.1 neurons labeled stochastically with different colors using Multi-Color Flip-Out (MCFO). Scale bar = 50 µm. **h**, Comparison of the skeletonized light microscopy (LM) image of FB6-Split 2 and the EM skeleton from the Hemibrain dataset.

To further analyze the morphology of the two single-neuron split-GAL4 lines, we performed multi-color flip-out (MCFO, Nern et al., 2015) enabling us to stochastically label individual neurons and hence giving us access to their fine anatomy (Fig. 4g). These experiments revealed a class of neurons with characteristic branching in the SMP and FB layer 6 and large soma (Fig. 4g, white arrows). To identify the cell type identity of these neurons, we morphologically compared light microscopy (LM) images to the FB6 cell types in the hemibrain electron microscopy (EM) connectome dataset (version 1.2.1; Scheffer et al., 2020; Hulse et al., 2021) using computational neuroanatomical tools (Bates et al., 2020; Suppl. Fig. 14). Using NBLAST scoring (Costa et al., 2016) as a similarity metric and visual inspection of the neuronal morphologies, we found that our Split-GAL4 driver lines very likely label a FB6A neuron (Fig. 4h and Suppl. Fig. 14d and e). While we consistently only observed one FB6A neuron labeled by either of the two FB6 Split-GAL4 driver lines, the hemibrain dataset identified 3 FB6A neurons per hemisphere with indistinguishable morphology and similar connectivity patterns in the SNP and FB6. This suggests that these three anatomically very similar neurons may not be identical at the genetic, and therefore functional level. For this reason we will refer to the cells we identified in our SplitGAL4 lines as FB6A.1. We have therefore identified a subset of neurons projecting to the SNP and FB6 that are important for regulating the probability of the fly transitioning from exploration to exploitation. While our neurogenetic dissection supports the importance of the FB6A.1 cell type in mediating this computation, complex foraging decisions almost certainly require computations performed by a larger neuronal network.

### Neuronal activity in FB6A.1 neurons is modulated by protein state

Which computations might necessitate FB6A.1 modulation of the probability to transition from exploration to exploitation? Within foraging theory, the budget theory predicts that foraging parameters can be dependent on the internal nutritional state of the animal (Stephens, 1981; Caraco et al., 1980; Kacelnik and El Mouden, 2013). Indeed, internal states such as hunger, reproductive state, and the lack of specific nutrients have a profound impact on animal behavior (Münch et al., 2020). Accordingly, previous studies have shown that sleep state and the fly’s energy state affect activity in the CX, including in the FB (Kempf et al., 2019). Given our results that the protein state of the animal alters its probability to engage with a food patch (Corrales-Carvajal et al., 2016 and Fig. 1d), we next examined whether activity in FB6A.1 neurons is altered by protein deprivation. To test this possibility, we performed whole-cell patch-clamp recordings of FB6A.1 neurons in either fully fed or protein-deprived flies (Fig. 5a-c). Strikingly, deprived flies exhibited significantly lower firing rates (Fig. 5b-d) and resting membrane potentials (Fig. 5e), showing that FB6A.1 neuron activity is modulated by the fly’s protein state. The decreased firing rate and resting membrane potential after protein deprivation are also consistent with the FB6A.1 silencing phenotype (Fig. 4c and e), where the probability to engage with a yeast patch is increased upon silencing. Together, these findings suggest that protein state alters FB6A.1 activity to modulate the probability to transition from exploration to exploitation of a yeast patch.

**Figure 5.**
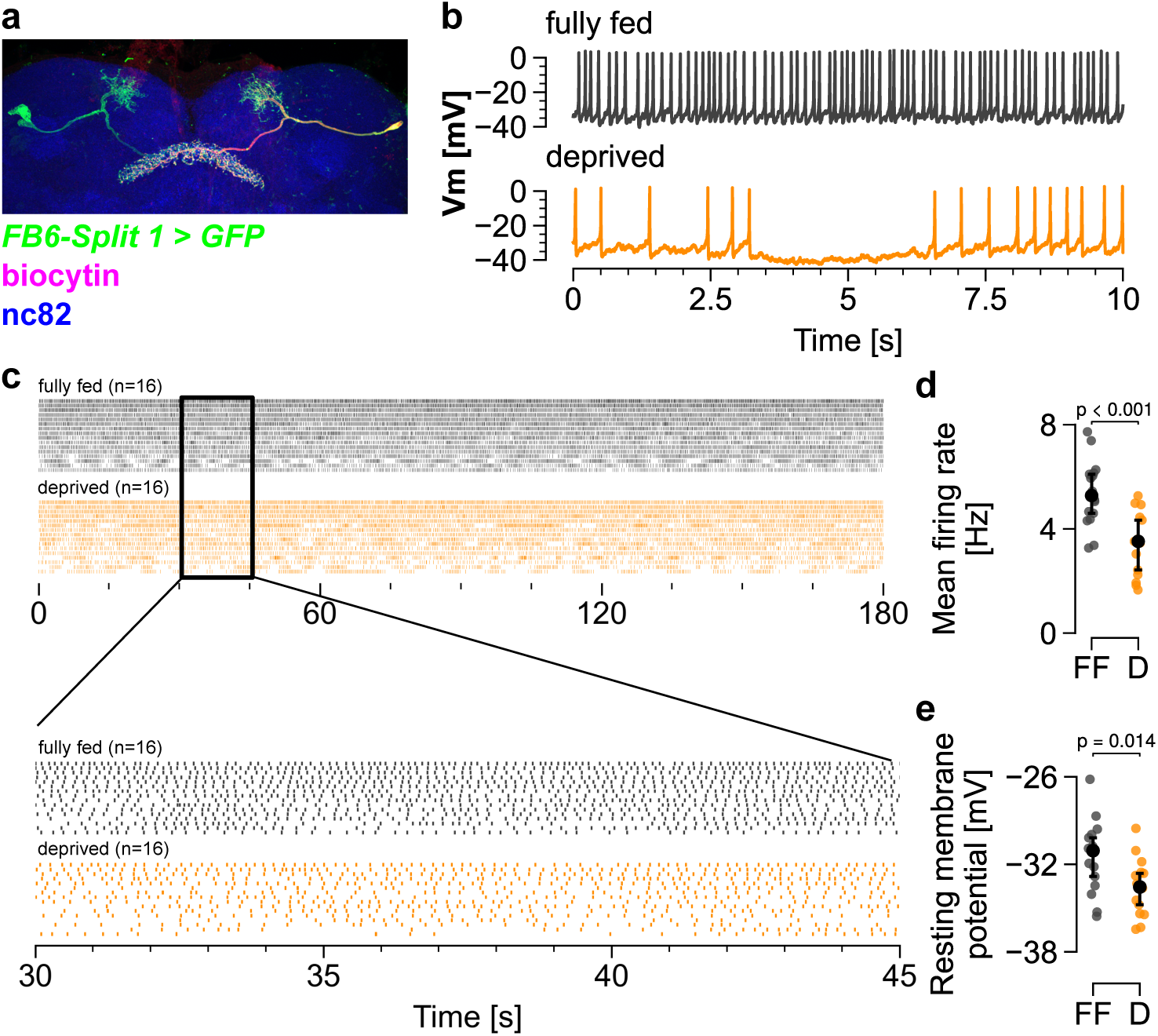
**Activity of the FB6A.1 neuron is modulated by the protein state of the fly. a**, Maximum intensity projection of a confocal stack of FB6A.1 neurons labeled by *FB6-Split 1*-driven expression of GFP as well as biocytin filling after whole-cell patch-clamp recording. **b**, Examples of membrane potential (Vm [mV]) traces for fully fed and protein deprived flies. **c**, Spike raster plots for fully fed and protein deprived flies recorded for three minutes. Inset shows 15 s of recordings. n = 16. **d**, Mean firing rate calculated over the whole recording shown in **(c)**. **e**, Mean resting membrane potential calculated on filtered, sub-threshold signal. **d-e**, P-values are obtained by performing Wilcoxon rank-sum tests, *n =* 16.

### Flies adapt foraging decisions to the density of food patches

A classic prediction of patch foraging models is that patch density—and therefore the time traveled between food patches—should influence different decision parameters, especially how long the animal exploits a food patch (Stephens and Krebs, 1986). One of the most prominent quantitative formalizations of this idea is the Marginal Value Theorem (MVT), which suggests that the decision of when to leave a patch is determined by the comparison between the cost of transitioning and exploiting other patches, and the current rate of energy gain at the patch (Charnov, 1976). The intuition behind these theories is that an increase in the distance between food patches increases the cost of transitioning, which should lead to an increase in the value of the food patches, and hence how long an animal exploits it. This relationship has been widely reproduced in animals foraging in the wild as well as in more reduced rodent and human experimental settings (Kacelnik, 1984; Kane et al., 2019; Kolling et al., 2012). To test if the food patch density (that is, the time required to transition between food patches) also alters foraging decisions in Drosophila, we designed a larger foraging arena in which the food patch geometry and content were the same as those used in all previously described experiments with the only difference being a significantly larger distance between food patches (Fig. 6a). Indeed, we found that the travel time between patches was significantly shorter in the arena with higher patch density (Suppl. Fig. 15). Classical patch foraging models, including MVT, predict that flies foraging in the large arena should have a higher mean duration of yeast visits. Indeed, flies foraging in the large arena (low patch density) showed a significantly higher mean duration of yeast visits when compared to flies foraging in the small arena (high patch density) (Fig. 6b). Importantly, feeding behavior, as measured using total micromovements on yeast, did not differ between the two arenas, highlighting the specificity of the effect (Suppl. Fig. 16). Most literature on the relationship between patch density and foraging decisions focuses on the time spent on a patch. To test if patch density also alters the probability to transition from exploration to exploitation, we compared the probability to engage with yeast patches between flies foraging in the small or large arena. Flies foraging in the environment with a low patch density had a significantly higher probability to engage with yeast patches (Fig. 6c). These experiments show that Drosophila adapts its foraging decisions to the density of the food in the environment. It does so in a way that follows the predictions made by classic foraging theory, highlighting the applicability of patch foraging models to a wide range of organisms, including *Drosophila*. Finally, our data demonstrate the ability of invertebrates to adapt their behavior according to statistical features of their environment, such as the density of food patches.

**Figure 6.**
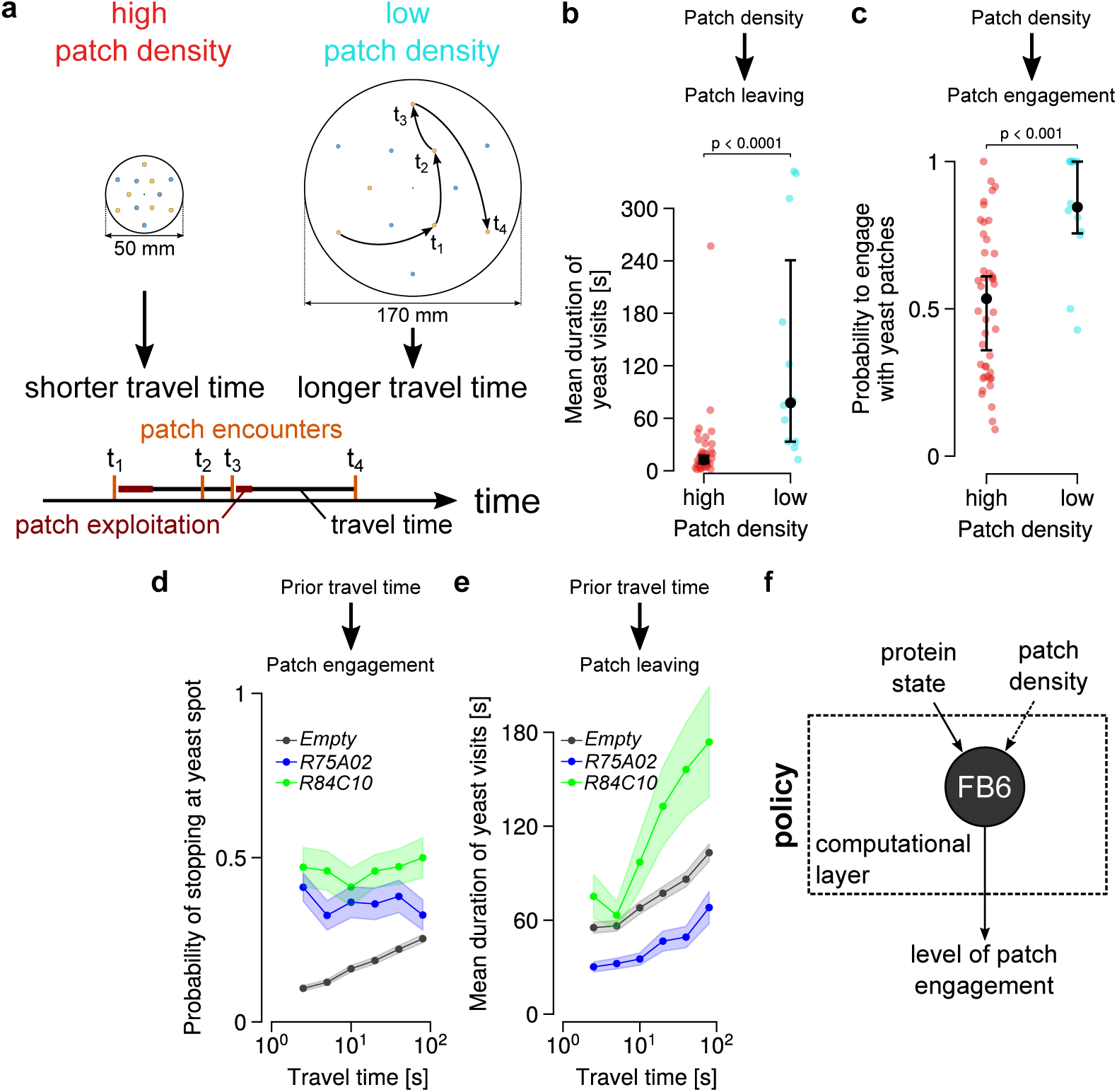
**Flies adapt foraging decisions to the patch density in the foraging environment. a**, Diagram of the two foraging arenas of different sizes used in the patch density experiments. The different distances between patches result in an either high or low patch density and corresponding shorter and longer travel times, respectively (top). Schematic of the logic used to calculate travel times (bottom). **b**, Mean duration of yeast visits comparing flies foraging in the high or low patch density environments. **c**, Probability to engage with yeast patches comparing flies foraging in the high or and low patch density environments, *n =* 12-44. **d**, Effect of silencing neurons labeled by *R84C10-* and *R75A02-GAL4* on the probability to engage with yeast patches across binned travel times (relationship between the probability to engage and the travel time between patches). **e**, Effect of silencing neurons labeled by *R84C10-* and *R75A02-GAL4* on the mean duration of yeast visits across binned travel times (relationship between the mean duration of a visit and the travel time between patches). For **d** and **e**, the n for the different experiments are: experimental *n =* 45-52. control *n =* 361. Dot and error bars in (**b-e**) indicate medians and IQR, data points correspond to individual flies. P-values are obtained by performing Wilcoxon rank sum tests. **f**, FB6 is part of a computational layer integrating state information to change policies regulating patch engagement.

### FB6 tangential neurons are important to specifically adapt patch engagement to the patch density of a given environment

How might flies estimate the density of food patches to alter their foraging decisions? A simple model of how they might do so follows from tenets of foraging theory, especially MVT: the cost of transitioning between patches should be correlated with travel time (Charnov, 1976). Therefore, the animal should adapt its foraging decisions (such as the time on the patch or the probability to engage) to the travel time between patch encounters. By doing so, different decision-making parameters will, on average, be adapted to the density of patches in the environment. To test if flies follow this simple rule, we analyzed how travel time relates to the probability to engage in the next visit (Fig. 6a). Indeed, control flies (*Empty* in gray) showed a monotonic increase in the probability to engage with increasing travel time (Fig. 6d). Given that FB6 neurons modulate the probability to engage, we tested if silencing FB6 neurons would affect how flies adapt foraging parameters based on travel distance. Silencing FB6 neurons using different GAL4 lines abolished the ability of flies to adapt the probability to engage in response to differences in the travel time between patches (Fig. 6d). Importantly, changing the protein state of the animal does not abolish the ability of the fly to adapt its foraging decisions to the travel time but simply shifts the relationship towards higher probabilities (Suppl. Fig. 17). This strongly suggests that the silencing phenotype cannot be explained by the manipulation only mimicking a protein-deprived state. To test if this phenotype is specific to the probability to engage, we tested if flies adapt how long they stay on a food patch according to their travel time. Indeed, as predicted by foraging theory, longer travel times lead to an increase in the time spent on a food patch (Fig. 6e). In contrast to the probability to engage, silencing FB6 neurons did not abolish the ability of the fly to adapt this foraging decision to the travel time. Thus, silencing FB6 neurons does not perturb the fly’s ability to estimate its travel time but rather its ability to translate that into a change in engagement probability. It also emphasizes the specificity of FB6 neurons in modulating the probability to engage, while leaving other foraging parameters unaltered. In conclusion, we identify a novel role for a specific subset of dorsal FB neurons in integrating internal state information (protein state) with external information about the statistics of the environment, to control a foraging decision which defines the trade-off between exploration and exploitation.

## Discussion

Foraging is an essential, highly complex, and ethologically relevant facet of animal behavior. Here, we used video-based tracking of individual flies and genetic manipulations of neuronal activity to dissect the neural underpinnings of a key strategic aspect of patch foraging: the exploration-exploitation trade-off. First, building on our previous findings (Corrales-Carvajal et al., 2016) we showed that this trade-off is tightly controlled by the nutritional state of the fly, where flies modulate foraging decisions to specifically compensate for a nutrient that they lack (that is, amino acids or carbohydrates). Then, we used a large-scale neuronal silencing screen followed by intersectional genetic refinement of the expression patterns of GAL4 lines showing phenotypes, to identify a bilateral pair of neurons in the dorsal fan-shaped body (FB6A.1) that control the probability to engage with a food patch. Strikingly, manipulations of the activity of FB6A.1 neurons did not affect other foraging decisions (for example, mean duration of visits on a food patch) or the cumulative food intake of the fly, suggesting that different foraging decisions are modularly controlled by distinct circuits. We furthermore found that protein deprivation decreased the activity of the FB6A.1 neuron. Importantly, the direction of this activity change is consistent with the observation that protein deprivation and FB6A.1 neuronal silencing produce similar behavioral phenotypes. Our findings reiterate the power of using large-scale, unbiased behavioral screens in *Drosophila* to identify mechanistic elements controlling complex behaviors and demonstrate how detailed behavioral analysis can identify state-modulated neurons controlling complex decisions in naturalistic settings.

How might FB6A.1 neurons alter the probability to engage, and hence modulate the transition from exploration to exploitation? Intriguingly, the dFB has emerged as a neuropil modulating multiple state-dependent behaviors (Donlea et al., 2014; Sareen et al., 2021; Musso et al., 2021). Most prominent is the role of dFB neurons in modulating sleep upon sleep deprivation and changes in diet (Donlea et al., 2014, 2018; French et al., 2021). These studies and our work suggest that dFB neurons receive internal state information (be it about the sleep state or the nutritional state of the animal) to gate how sensory information is used to shape motor outputs (Donlea et al., 2014, 2018; Carvalho-Santos and Ribeiro, 2023; Raccuglia et al., 2022). Therefore, we propose that FB6 neurons receive information about the internal protein state of the animal as well as the density of food in the environment, and modulate the animal’s decision on whether or not to engage with a food patch. Given the high degree of interconnectivity between different dFB neurons and other FB cell types (Hulse et al., 2021), it is likely that these foraging computations will involve complex interactions with other FB cell types in addition to FB6 neurons. As such, an attractive hypothesis is that FB6 neurons are part of a computational layer that implements the policy of how internal and external states map onto a specific set of actions, thereby shaping the decision-making strategies that define foraging behavior (Fig. 6f). Our work emphasizes the integrative role of the CX as a neuropil receiving different types of state and sensory information in order to guide behavioral strategies (Hulse et al., 2021; Dan et al., 2021).

Significant progress in our understanding of the neuronal basis of foraging computations has come from an integration of concepts from classical foraging theory, neuroeconomics, and systems neuroscience (Hayden et al., 2011; Kolling et al., 2012; Blanchard and Hayden, 2014; Lottem et al., 2018; Le Heron et al., 2020; Vertechi et al., 2020). Most studies use measurements of neuronal activity in rodents, primates or humans performing sophisticated but reduced behavioral tasks aimed at mimicking free foraging behavior, to identify neuronal populations whose activity correlates with quantitative predictions from theory (Hayden et al., 2011; Kolling et al., 2012; Lottem et al., 2018). These studies have mainly focused on the decision to leave a “patch” to search for new ones and have identified neurons in, for example, the ventromedial prefrontal cortex (vmPFC) and the anterior cingulate cortex (ACC) as encoding variables that contribute to this decision-making process (Hayden et al., 2011; Kolling et al., 2012, 2016; Le Heron et al., 2020). While aspects of these findings have generated some controversy (Kolling et al., 2016; Kane et al., 2022), these studies have begun to provide a way to think about how brains compute and optimize complex decisions. However, substantial questions remain. Most prior studies rely on extensive training of animals and it is not clear how well the reduced tasks employed mimic behavior in naturalistic settings. Furthermore, while specific brain regions have been identified as playing important roles in these computations, the architecture of the underlying circuits, the identity of the involved neurons, and the extent to which they are causally involved in shaping foraging decisions, is not known. Nor is it known how nutritional states modulate the underlying neuronal substrates to shape the required computations (Münch et al., 2020).

Our identification of a single neuron that influences the probability of transitioning from exploration to exploitation based on both the protein state of the animal and the density of food patches in the environment provides a new entry point to dissect these types of decisions. Using the neurogenetic approaches available in the fly, including recent progress in imaging neuronal activity and the availability of synaptic resolution connectomes, should enable progress in understanding how these complex decisions are made. Our data indicate that FB6 neurons might participate in computing foraging decisions according to principles which are related to the MVT. While more quantitative data are required to test if and how flies adapt foraging decisions to the statistics of the environment and how the activity of FB6 neurons might contribute to these computations, it is attractive to speculate that the CX and more precisely the neurons we identified in the dFB are part of the substrate that implements universal foraging computations as proposed by the MVT.

While complex foraging computations and decisionmaking strategies have been mainly documented in vertebrates (Shadlen and Kiani, 2013; Cazettes et al., 2023) and social insects (Deneubourg and Goss, 1989; Greggers and Menzel, 1993; Montague et al., 1995; Chittka et al., 2003), a growing literature indicates that other animals with a numerically simple nervous system such as nematodes and flies also perform such computations (Calhoun et al., 2015; Flavell et al., 2013; Wosniack et al., 2022; Katzen et al., 2023; Vijayan et al., 2023). To what extent these animals can infer and use statistical properties of the environment to perform neuronal computations remains controversial. Here we show that Drosophila adapt foraging decisions according to the density of food patches in their environment. We also provide evidence that they do so by matching the decisions to the travel time between patches and that this process requires activity in FB6 neurons. Given the importance of nutrients in shaping the fitness of all organisms, it might seem obvious that all animals should have evolved the capacity to optimize foraging and that this capacity could have reached a relatively high level of sophistication. As such, foraging is likely to have served as an ancient blueprint for the evolution of more complex cognitive abilities. Comparing how different brains implement foraging computations will give us valuable insights into how cognition might have evolved.

Our findings highlight the profound impact nutritional states have on complex behaviors and how the combination of quantitative and high-throughput behavioral screens with neurogenetic and neuroanatomical approaches can reveal the neuronal substrate for this regulation. By what mechanisms might protein state affect the probability to engage? Given that other protein-state sensitive foraging decisions are not affected by FB6A.1 manipulations, it is possible that FB6A.1 neurons do not directly sense the protein state of the animal. Instead, protein state could be relayed by neuromodulators secreted by protein-sensitive neurons which broadcast the nutritional state of the animal more broadly, as has been proposed for other nutrient-sensitive behaviors (Sareen et al., 2021; Musso et al., 2021). Identifying these neurons and mapping out how they affect FB6A.1 activity will be an important next step in understanding the molecular and cellular mechanisms by which internal states shape the exploration-exploitation tradeoff and therefore how risk is computed and managed at the level of neuronal circuits. Taken together our work highlights the ability of invertebrates to adapt complex foraging decisions according to internal and external information, pinpoints the dFB, and, more generally, the CX, a highly conserved brain structure in insects as a central brain region for performing these complex computations. Combining anatomical information at synaptic resolution (Hulse et al., 2021) with quantitative foraging theories (Stephens and Krebs, 1986; Davidson and El Hady, 2019), sophisticated behavioral paradigms (Haberkern et al., 2019; Matheson et al., 2022), and transgenic lines that provide cell-specific access to CX neurons (Wolff et al., 2015; Wolff and Rubin, 2018) will open the way to a deeper understanding of how foraging computations are implemented at the level of defined neuron types. Our work therefore provides an entry point into a computational and neuron-level understanding of how brains optimize foraging decisions to maintain nutrient homeostasis.

## ACKNOWLEDGEMENTS

We thank A.P. Elias for technical assistance with the fly brain dissections and immunostainings; A. Laborde for technical assistance with building the PC infrastructure for the tracking data acquisition; the Champalimaud Hardware and Software Platform, specifically A. Silva, P. Carriço, D. Bento and F. Carvalho for building hardware and software of the LED boards used for closed-loop optogenetics; the Champalimaud Fly Platform for support with fly brain dissections and immunostainings; the Champalimaud Glass Wash and Media Preparation Platform for help with cleaning arenas for the silencing screen; J. Schweizer for help with flyPAD experiments; the Janelia Project Technical Resources team (led by G. Ihrke) performed electrophysiology experiments (J-Y.P.), brain dissections and staining (K. Close), and confocal imaging (C. Christoforou); E. Chiappe, L. Vasconcelos, R. Benton, D. Anderson and N. Yapici for sharing fly lines; R. Henriques for creating the LAT_E_X template. A.M. Hermundstad, E. Chiappe, C. Machens, Z. Santos-Carvalho, S.J. Walker, S. Henriques, R. Beresford and the whole Behaviour and Metabolism laboratory for many fruitful discussions, valuable feedback throughout the project, and comments on the manuscript. Fly lines were obtained from the Bloomington Drosophila Stock Center (NIH P40OD018537) and the Janelia Fly Facility. We are grateful to the Janelia Visitor Scientist Program for their support of the project. D.G. was supported by a doctoral fellowship PD/BD/114273/2016 from the Portuguese Foundation for Science and Technology (FCT). I.T. was supported by a Marie Skłodowska-Curie postdoctoral fellowship (H2020-WF-01-2018-867459 to Ibrahim Tastekin), and FCT–Fundação para a Ciência e Tecnologia, grant PTDC/MEDNEU/4001/2021. D.M. was supported by a DFG Research Fellowship (MU 4116/1-1). The project leading to these results has received funding from the grant PTDC/MED-NEU/4001/2021 from the Portuguese Foundation for Science and Technology (FCT) awarded to C.R.. The project was supported by the research infrastructure Congento, co-financed by Lisboa Regional Operational Programme (Lisboa2020), under the PORTUGAL 2020 Partnership Agreement, through the European Regional Development Fund (ERDF) and FCT–Fundação para a Ciência e Tecnologia (Portugal) under the project LISBOA-010145-FEDER-022170. Research at Janelia is supported by the Howard Hughes Medical Institute, which also supported H.H., V.J., J-Y.P. and G.M.R.. Research at the Centre for the Unknown is supported by the Champalimaud Foundation.

## Materials and Methods

### Fly husbandry

Flies were reared on yeast-based medium (per liter of water: 8 g agar (NZYTech, Portugal), 80 g barley malt syrup (Próvida, Portugal), 22 g sugar beet syrup (Grafschafter, Germany), 80 g cornflour (Próvida, Portugal), 10 g soya flour (A. Centazi, Portugal), 18 g instant yeast (Saf-instant, Lesaffre, France), 8 mL propionic acid (Argos), and 12 mL nipagin (Tegosept, Dutscher, France) (15% in 96% ethanol) supplemented with instant yeast granules on the surface (Saf-instant, Lesaffre, France)) and kept at 25*^◦^* C, 70% relative humidity in a 12 h/ 12 h light-dark cycle.

### Internal state manipulations and holidic media

Holidic media for amino acid (AA) and carbohydrate deprivation were prepared as described previously (Leitão-Gonçalves et al., 2017) using the exome-matched formulation with food preservatives (Piper et al., 2017). For AA-deprived flies, all amino acids contained in the medium were not added to the diet, while keeping the concentration of all other nutrients constant. Likewise, for carbohydrate-deprived flies, all carbohydrates were removed from the medium. For all experiments using the holidic medium, the following dietary treatment protocol was used to ensure a fully-fed state: groups of 0*±*4-day old flies (10 females) were collected into fresh YBM-filled vials, 5 Canton-S males were added to the vials to ensure females were mated, and transferred to fresh YBM after 48 hours. After 24 hours, flies were transferred to different holidic media and kept there for 3 days. Protein deprivation was induced by transferring 2-6 days old, mated female flies to vials containing a paper tissue soaked with 5 mL of 100 mM sucrose solution for 5 days. Fully fed flies were age-matched with protein-deprived flies. All flies were transferred to vials with the respective fresh medium every two days. Mating was ensured by keeping 10 female flies together with 5 male flies.

### Fly stocks and genetics

The list of stocks used in this study are shown in Supplementary Table 1 below.

Detailed fly genotypes used were:

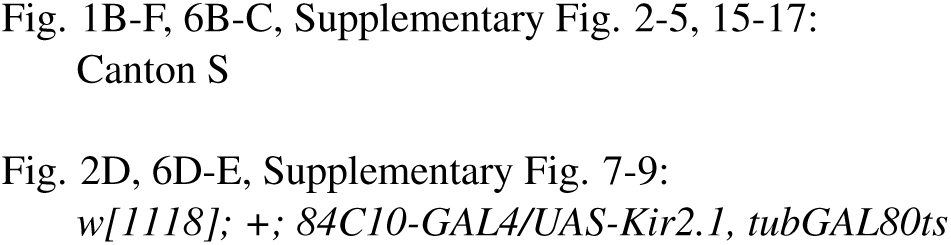
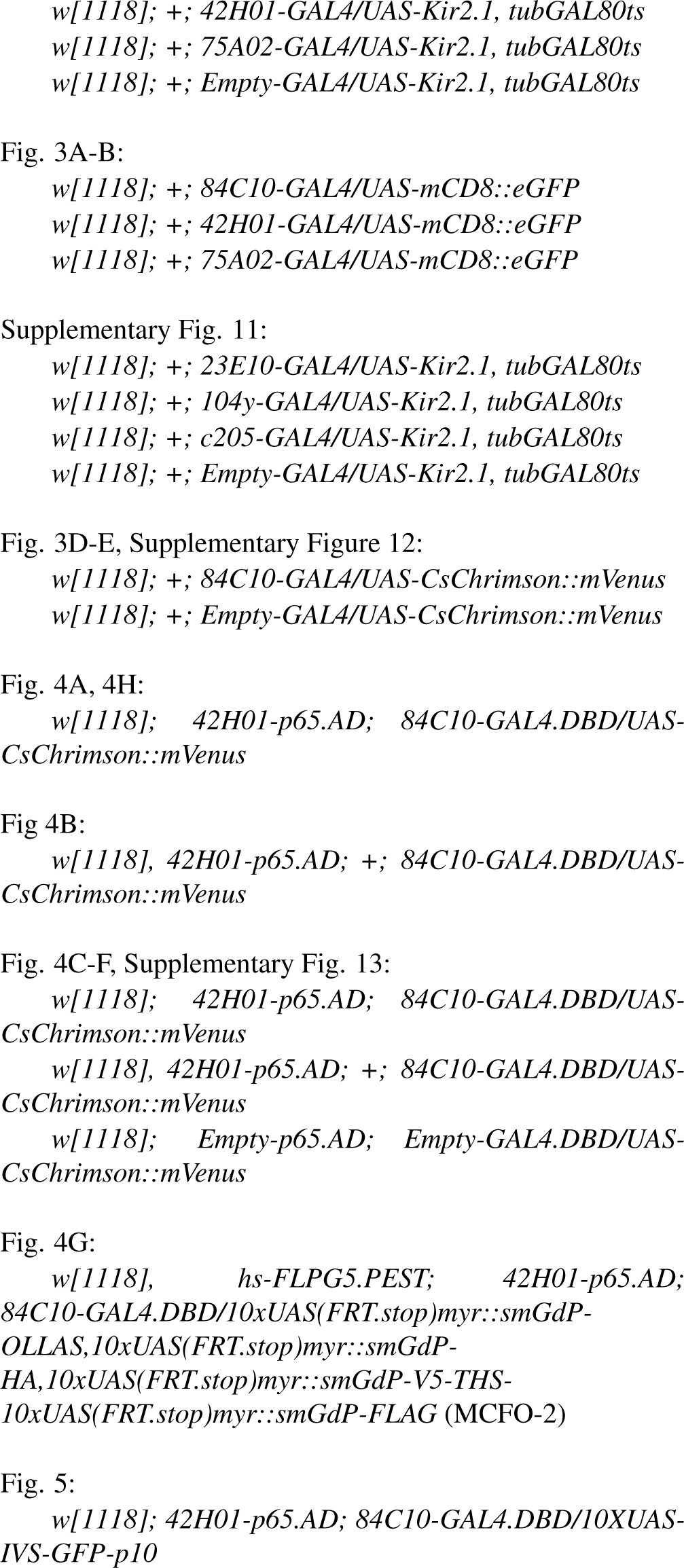

### Split-GAL4 lines

Split-GAL4 lines were constructed and characterized by the Rubin lab as in previous work (Dionne et al., 2018). Briefly, we identified candidate AD and DBD hemidriver pairs (Tirian and Dickson, 2017; Dionne et al., 2018) for expression in cell types of interest and assembled successful combinations into stable fly strains that were then used for subsequent experiments. All Split-GAL4 drivers created in this study are listed in Supplementary Table 1. For testing the Split-GAL4 lines, we used an *Empty Split-GAL4* driver line (BDSC #79603) as control.

### Foraging assays

The standard foraging assay is performed in an arena containing 5 µL patches of 10% yeast and 100 mM sucrose, respectively, distributed in a radial pattern (see Fig. 1a, Suppl. Fig. 1 and section *Image-based tracking setup*). Per setup, we recorded fly trajectories of four individual flies in separated arenas for 1 hour. We built 12 of these setups which were connected to four PCs for simultaneous data acquisition. We used custom-written software for the data acquisition (see details in the subsection *Tracking algorithm*).

### Image-based tracking setup

We built a high-throughput tracking setup that can record four flies in individual arenas using a custom-built acrylic arena holder. Arenas were placed above an IR backlight for video recording using a FLIR Flea3 camera (FL3-U3-32S2M-CS, Teledyne FLIR LLC, USA) acquiring videos at 30 frames per second at a resolution of 1408 px *×* 1408 px. This resolution resulted in an average realworld pixel resolution of 7.8 px per mm.

### Tracking algorithm

We used a custom, real-time tracking algorithm using the Bonsai framework (Lopes et al., 2015). Fly centroids and trajectories were extracted using a background subtraction method, and for each recorded frame we measured the xand y-position of the centroid, major and minor axis as well as the angle of the centroid. We also developed custom-written Python scripts to determine the head position of the fly using an offline algorithm. This algorithm compares the pixel intensity of both extreme points of the centroid along the major axis to determine the head, which is darker than the tail position due to the transparency of the wings. We then propagated the head position by applying a proximity rule (Gomez-Marin et al., 2011), in which the position closer to the previous head position is assigned as head position. This rule becomes invalid when flies jump, therefore we detected jumps as segments to re-apply the pixel intensity comparison. We further validated correct head position assignment, by comparing the movement vector to the heading vector, which should largely align because flies mostly walk forwards. Using this head detection method, we validated the accuracy by manual visual inspection to be correct in more than 98% of frames (1000 frames randomly sampled).

### Behavioral classification

From the head and body trajectories, we calculated the displacements between each of the consecutive frames. By dividing these displacements by the time interval between the frames, we obtain the translational speeds of head and centroid positions. We subsequently filtered the speed time series using a Gaussian kernel with a window length of 36 frames and a standard deviation of 3.6. Similarly, we calculated the rotational speed from displacements in the angle of the centroid to head relative to the reference frame. Based on these kinematic variables, we used threshold values to classify behavioral segments. “Walking” segments are defined by instances, in which the fly’s head speed is above 2 mm/s, while “Jumps” are above 20 mm/s.

“Turns” are defined by instances, in which the fly’s rotational speed is larger than 125 *^◦^*/s. “Resting” segments are instances, in which the fly’s head speed is lower than 0.2 mm/s. “Micromovements” are instances, which have not been so far classified otherwise, and as such the head speed falls into the range of 0.2 to 2 mm/s. Micromovements which are performed closer than 2.5 mm to a yeast patch are considered “yeast micromovements”, while those closer than 2.5 mm to a sucrose patch are called “sucrose micromovements”. We filtered out spurious high body speeds that rarely occur during micromovements.

### Patch encounters, visits, and ratio of returns

“Encounters” are segments in which the fly is closer than 3 mm to a given food patch, regardless of whether feeding occurs or not. “Visits” are all micromovements that occur on the same food patch without the fly leaving a distance larger than 5 mm from the patch. The “probability to engage with a food patch” is calculated by dividing the total number of visits by the total number of encounters. Since each encounter can at most have one visit, the resulting ratio is between 0 and 1, and therefore can be described as a probability. For patch leaving decisions, we calculated the “mean duration of visits” for all patches of each substrate. The “ratio of returns to the same food patch” is calculated by dividing the number of returns to the same patch by the total number of visits to all patches of the same substrate.

### GAL4 silencing screen

We selected GAL4 driver lines with sparse expression in specific neuropils so that the whole collection covers a wide range of neuropils (see Suppl. Fig. 6). We expressed *Kir2.1* in different neuronal subsets using the GAL4 driver lines and used our high-throughput foraging assay to screen 434 GAL4 lines for phenotypes in the probability to engage as described above. We used a *tubGal80ts* transgene to induce the expression of Kir2.1 protein 24-48 hours before the assay, by moving the flies to the permissive temperature of 30 *^◦^*C. This way, we aimed to avoid any developmental effects that might be caused by the constitutive expression of Kir2.1. Briefly, we organized crosses, sorting and assays in a weekly schedule: As such, we sorted flies two weeks after setting up crosses, and performed the assay one week after sorting. The sorted flies were 0-4 days old and kept together with male Canton S flies to ensure mating. After one day on standard YBM, we flipped these flies to a new vial of standard YBM. After another day on this food, we protein deprived flies for 5 days as described above. On the day of the assays, we moved the flies to 25 *^◦^*C 1-2 hours before testing. Each experimental day of the screen consisted of testing 6-9 experimental lines together with the control flies (*Empty-GAL4* (BDSC #68384) crossed to *UASKir2.1,tub-Gal80ts*).

### flyPAD assays

flyPAD assays were performed as previously described (Itskov et al., 2014). Flies were given a choice between 10% yeast and 20 mM sucrose, both in a 1% agarose solution. Flies were individually transferred to flyPAD arenas by mouth aspiration and allowed to feed for 1 h at 25 *^◦^*C, 70% RH. flyPAD data were acquired using the Bonsai framework (Lopes et al., 2015), and analyzed in MATLAB (MATLAB version: 8.2 (R2013b), Natick, Massachusetts: The MathWorks Inc.; 2022) using custom-written software, as described before (Itskov et al., 2014).

### Optogenetic assays

We performed optogenetic assays using custom-built RGB/IR LED boards containing 9 highpower (10 W) RGB LEDs (ref. no. LZ4-00MA00; LED Engin, USA) per arena for optogenetic stimulation, and 16 infrared LEDs (ref. no. AA3528F3S, Kingbright, USA) for backlight illumination. Fully-fed flies were reared on YBM laced with 0.4 mM all-trans-retinal (#R2500, Sigma-Aldrich) made from a stock solution of 100 mM all-trans-retinal dissolved in ethanol. For protein deprivation, flies were transferred for 3 days to vials containing a piece of tissue paper soaked with 5 mL of 100 mM sucrose solution laced with 0.4 mM all-trans-retinal.

Closed-loop optogenetic assays using red LEDs at 623 nm were performed with two stimulation conditions: either the LED turns on when the fly approaches the food patch (10 mm distance) (in 90% of the cases), or the LED turns on after the fly has initiated feeding by using a shorter activation radius (2 mm) and a delay of 3 s to ensure that the fly engages in feeding (catch trials). We wrote custom software in the Bonsai framework to automatically control LED stimulation and switch between these two conditions probabilistically (Fig. 3c). The LED stimulation power density was adjusted to 5 mW/cm^2^.

### Travel time analysis

We calculated the travel time for each resource encounter as the time duration from the end of the last encounter of a patch of the same substrate, regardless of whether a visit occurs, to the beginning of the current encounter. We binned the observed travel times to calculate the probability to engage with food patches. The binning of travel times was determined by logarithmic scaling with overlaps to ensure a minimum number of data samples across all bins (2.5-25, 5-50, 10-100, 20-200, 40-400, 80-800, 160-1600 s). Revisits were removed from the calculation of the probability to engage with food patches to avoid bias for short travel times.

### Fluorescent staining and confocal imaging

Males from each GAL4 line were crossed to females homozygous for the fluorescent reporter lines. Three- to ten-day-old adult, fully-fed mated females carrying both the Gal4 driver and the UAS reporter were dissected. Samples were dissected in 4 *^◦^*C 1X PBS (10173433, Fisher Scientific, UK) after a quick passage through EtOH (4146052, Carlo Elba), and were then transferred to formaldehyde solution (4% paraformaldehyde, P6148, Sigma-Aldrich in 1X PBS + 10% Triton-X, X100, Sigma-Aldrich) and incubated for 20-30 min at room temperature. Samples were then washed three times in PBST (0.5% Triton-X in PBS) and then blocked with 10% normal goat serum (16210-064, Invitrogen) in PBST for 15-60 min at room temperature. Samples were then incubated in primary antibody solutions (Rabbit anti-GFP, TP401, Torrey Pines Biolabs at 1:6000 and Mouse anti-NC82, Developmental Studies Hybridoma Bank at 1:10 in 5% normal goat serum in PBST). Primary antibody incubations were performed for 3 days at 4 *^◦^*C with rocking. They were then washed in PBST 2-3 times for 10-15 min at room temperature. The secondary antibodies were applied (Anti-mouse A594, A11032, Invitrogen at 1:500 and Anti-rabbit A488, A11008, Invitrogen at 1:500 in 5% normal goat serum in PBST) and brains were then incubated for 3 days at 4 *^◦^*C with rocking. They were again washed in PBST 2-3 times for 10-15 min at room temperature. Samples were mounted in Vectashield Mounting Medium (H-1000, Vector Laboratories). Images were captured on an inverted Zeiss LSM 980 (Carl Zeiss Co., Oberkochen, Germany) using a Plan-ApoChromat 20x/0.8 air lens objective (Carl Zeiss Co., Oberkochen, Germany).

### Alignment to the Janelia standard brain

Confocal stacks were aligned to a standard reference template (JFRC2, Jenett et al., 2012) using a custom-written Bash script using the software ANTs (Avants et al., 2011).

### EM morphological analysis and cell-type matching across modalities

First, we converted the aligned confocal images of the neurons of interest into a skeleton using Vaa3d (Peng et al., 2014). Second, this skeleton was transformed into the JFRC2 reference space and converted into a vectorized point cloud using the Natverse package (Bates et al., 2020) in R (R Core Team, 2021). Then, we used the NBLAST algorithm (Costa et al., 2016) to calculate similarity scores between the skeletonized LM neuron and available layer 6 FB neurons from the hemibrain dataset using the NeuPrint interface (v1.2.1, Clements et al., 2020).

### Electrophysiology

FB6A.1 recordings were obtained from 3to 4-day old female flies and performed as described previously (Dombrovski et al., 2023). Flies were anesthetized on a peltier-driven cold plate, transferred to a chilled vacuum holder, and mounted to a customized recording plate. To reduce brain motion, the proboscis was fixed to the head by applying UV cure glue. To access the FB6A.1 soma, the posterior part of the cuticle, fat and trachea were removed using a syringe needle and fine forceps. The brain was continuously perfused with an extracellular saline containing (in mM): 103 NaCl, 3 KCl, 5 N-Tris (hydroxymethyl) methyl-2aminoethane-sulfonic acid, 8 trehalose, 10 glucose, 26 NaHCO3, 1 NaH2PO4, 1.5 CaCl2 and 4 MgCl2. Osmolarity was adjusted to 275 mOsm and the saline was bubbled with 95% O2 / 5% CO2 during the experiment. To disrupt the perineural sheath around the soma, collagenase (0.5 mg/ml) was applied locally and then a small amount of tissue was removed to gain patch pipette access. Patchclamp electrodes (5-7 M) were filled with a solution containing 140 potassium aspartate, 10 4-(2-hydroxyethyl)-1piperazineethanesulfonic acid, 1 ethylene glycol tetraacetic acid, 4 MgATP, 0.5 Na3GTP and 1 KCl. The pH was 7.3 and the osmolarity was adjusted to approximately 265 mOsm. To enable visualization of the patched neurons, the intracellular solution contained 13 mM biocytin hydrazide. Recordings were acquired in current-clamp mode on a MultiClamp 700B amplifier, low-pass filtered at 10 kHz, digitized at 40 kHz, and analyzed using Clampfit 11 software (Molecular Devices). Resting membrane potential was estimated by smoothing the membrane potentials (Butterworth filter at 5 Hz).

### Immunohistochemistry of electrophysiological experiments

To visualize biocytin-filled neurons, the dissections, immunohistochemistry, and imaging of fly brains were done principally as previously described (Aso et al., 2014; https://www.janelia.org/project-team/flylight/protocols). In brief, brains were dissected in Schneider’s insect medium (omitting the ethanol rinse) and fixed in 2% paraformaldehyde (diluted in the same medium) at room temperature for 55 min. Tissues were washed in PBT (0.5% Triton X-100 in phosphate buffered saline) and blocked using 5% normal goat serum before incubation with antibodies and fluorescently labeled Streptavidin.

Tissues expressing GFP were stained with rabbit antiGFP (ThermoFisher Scientific A-11122, 1:1000) and mouse anti-BRP hybridoma supernatant (nc82, Developmental Studies Hybridoma Bank, Univ. Iowa, 1:30), followed by Alexa Fluor® 488-conjugated goat anti-rabbit and Alexa Fluor® 568-conjugated goat anti-mouse antibodies (ThermoFisher Scientific A-11034 and A-11031), respectively. Alexa Fluor® 594-conjugated streptavidin (ThermoFisher Scientific S-32356, 1:1000) was included with secondary antibody incubations. After staining and post-fixation in 4% paraformaldehyde, tissues were mounted on poly-L-lysine-coated cover slips, cleared, and embedded in DPX as described.

Image z-stacks were collected using an LSM880 confocal microscope (Zeiss, Germany) fitted with PlanApochromat 20x/0.8 M27 air and Plan-Apochromat 40x/1.3 M27 oil immersion objectives. The voxel size was 0.52 *×* 0.52 *×* 1.0 µm^3^ (1024 *×* 1024 pixels in xy) and 0.44 *×* 0.44 *×* 0.44 µm^3^ (688 *×* 688 pixels in xy) in 20x and 40x image stacks, respectively.

### Statistical analysis

For all statistical analyses, if not otherwise stated, we use non-parametric, two-sided hypothesis testing (i.e., Wilcoxon-Mann-Whitney rank sum tests). Specifically, we used the SciPy 1.8.0 (Virtanen et al., 2020) implementation of the function `stats.ranksum`.

### Data and code availability

We stored all source data and Python/R custom-written visualization scripts, which will be made available upon publication. Hardware and software files for the performing data acquisition and analysis will be made available upon publication.

## Author Contributions

Conceptualization, D.G., I.T., C.R.; Data Curation and formal analysis, D.G., I.T., D.M., J-Y.P., H.H., G.M.R., C.R.; Investigation, D.G., I.T., J-Y.P.. H.H., L.B., C.B., G.M.R., C.R.; Behavioral screen, D.G., I.T., D.M., L.S., C.B., H.H., G.M.R., C.R.: Electrophysiological recordings, J-Y.P.; Supervision, V.J., C.R.; Validation, D.G., I.T., J-Y.P., H.H., G.M.R., C.R.; Visualization, D.G., I.T., J-Y.P., H.H., G.M.R., C.R.; Writing – Original Draft, D.G., I.T., C.R.; Writing – Review & Editing, D.G., I.T., D.M., J-Y.P., H.H., V.J., G.M.R., C.R.; Project Administration, D.G., C.R.; Funding acquisition, V.J., G.M.R., C.R.. All authors read and approved the final manuscript.

## Supplementary materials

**Table 1.**
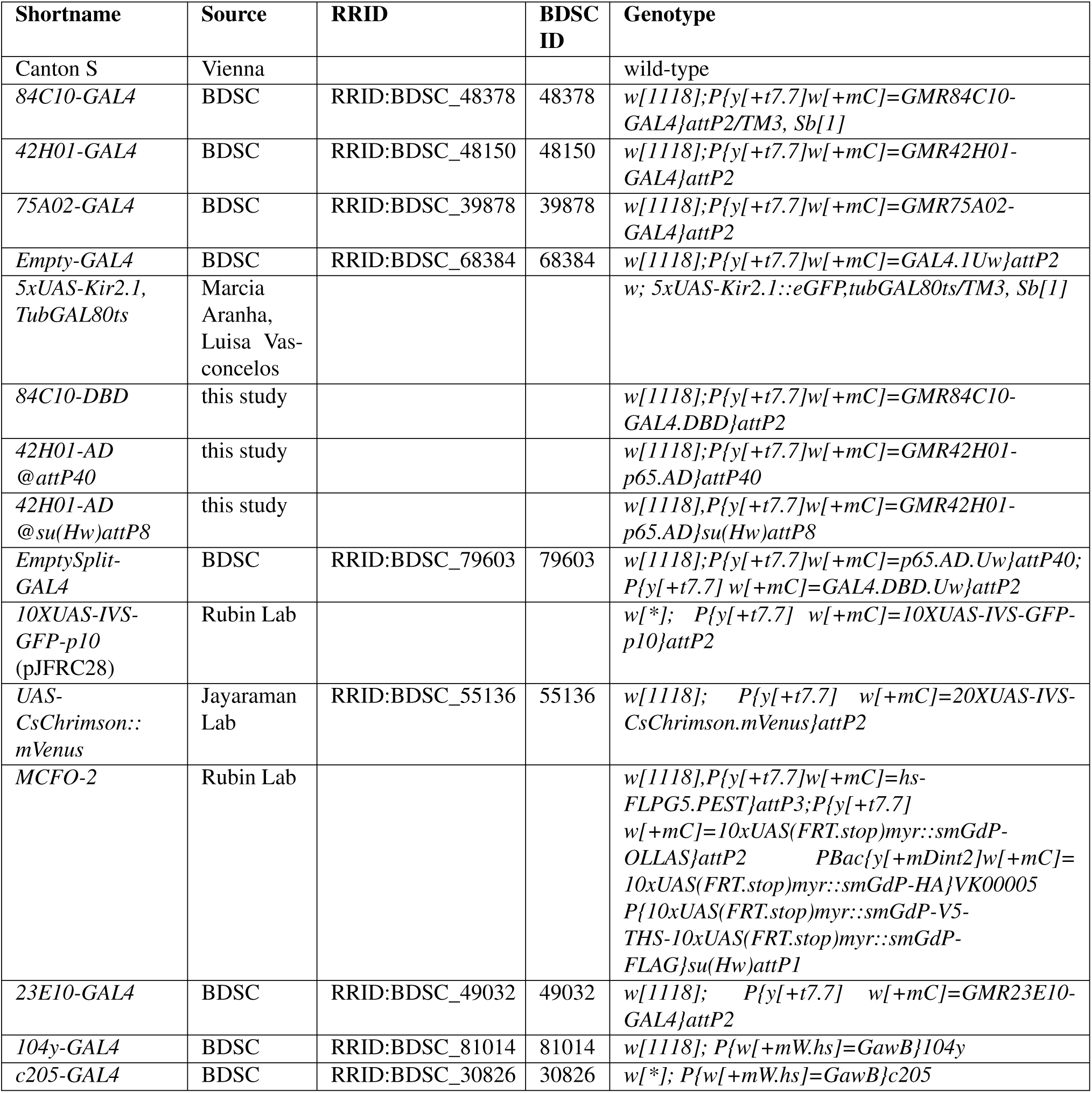
Stocks used in this study.

**Supplementary Figure 1.**
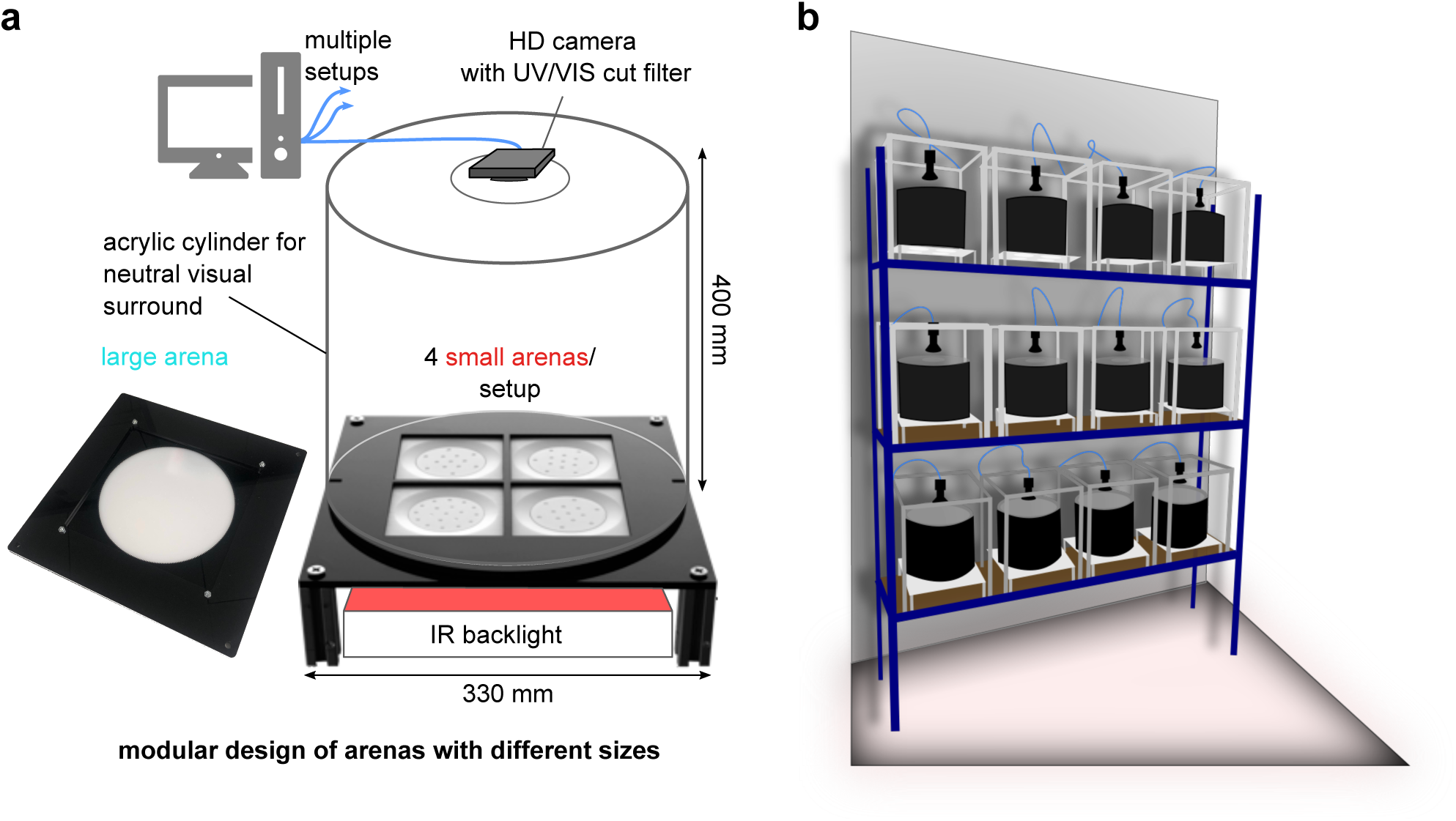
**High-throughput modular experimental framework for studying foraging decisions in flies. a**, Details of a single video tracking setup with the two different foraging arena designs (small and large). The small arena design was used in all experiments except when mentioned (for the patch density experiments). **b**, Schematic of the organization of the 12 modular setups used to perform the large-scale silencing screen in the temperature- and humidity-controlled behavior room.

**Supplementary Figure 2.**
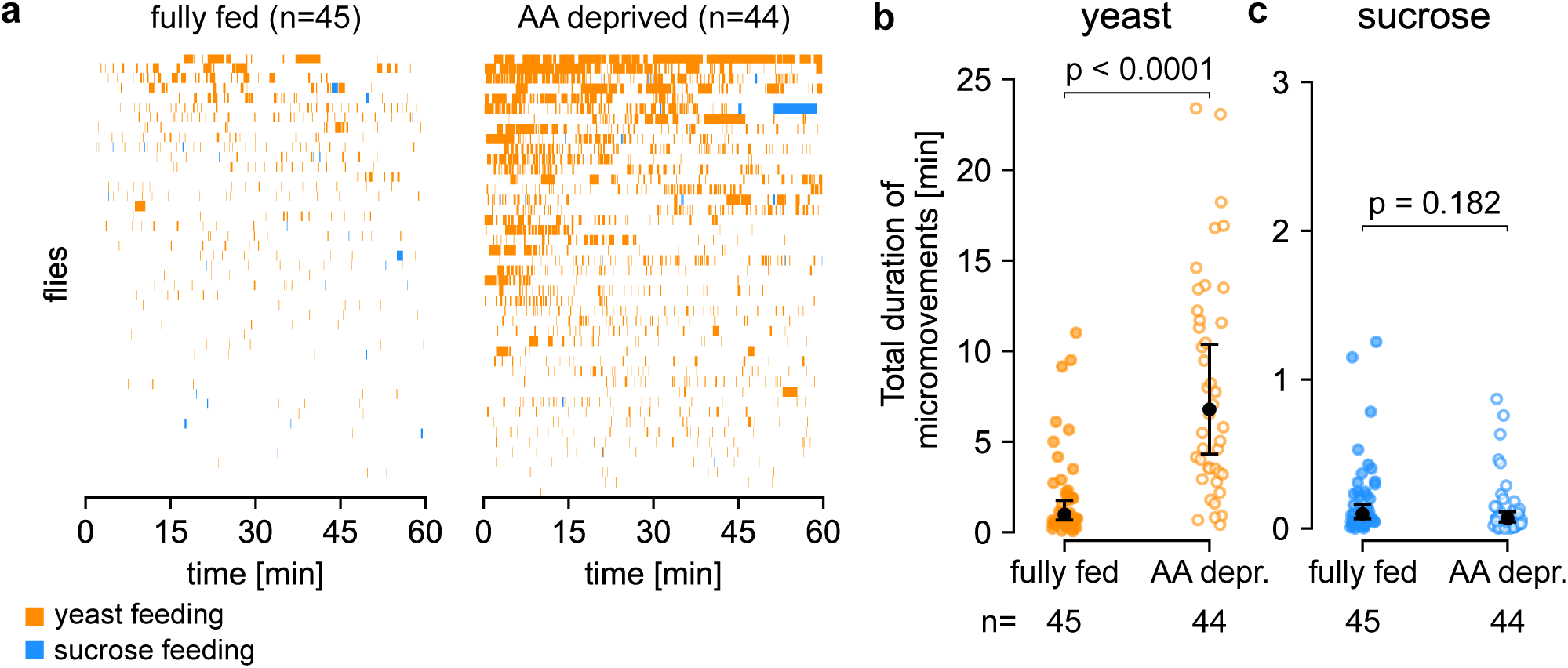
**Amino acid (AA) deprived flies increase the total amount of exploitation on yeast but not on sucrose patches. a**, Ethogram of yeast and sucrose feeding micromovements for fully fed and AA deprived flies. Each row represents the behavioral time course of one fly, and flies are sorted by the total duration of yeast micromovements. **b**, Total duration of yeast micromovements for fully fed and AA deprived flies. **c**, Total duration of sucrose micromovements for fully fed and AA deprived flies. **b-c**, Data points are individual flies, black dots and bars indicate median, and extrema within 1.5 of the inter-quartile range (IQR), respectively. P-values are obtained by performing Wilcoxon rank-sum tests, *n =* 44-45.

**Supplementary Figure 3.**
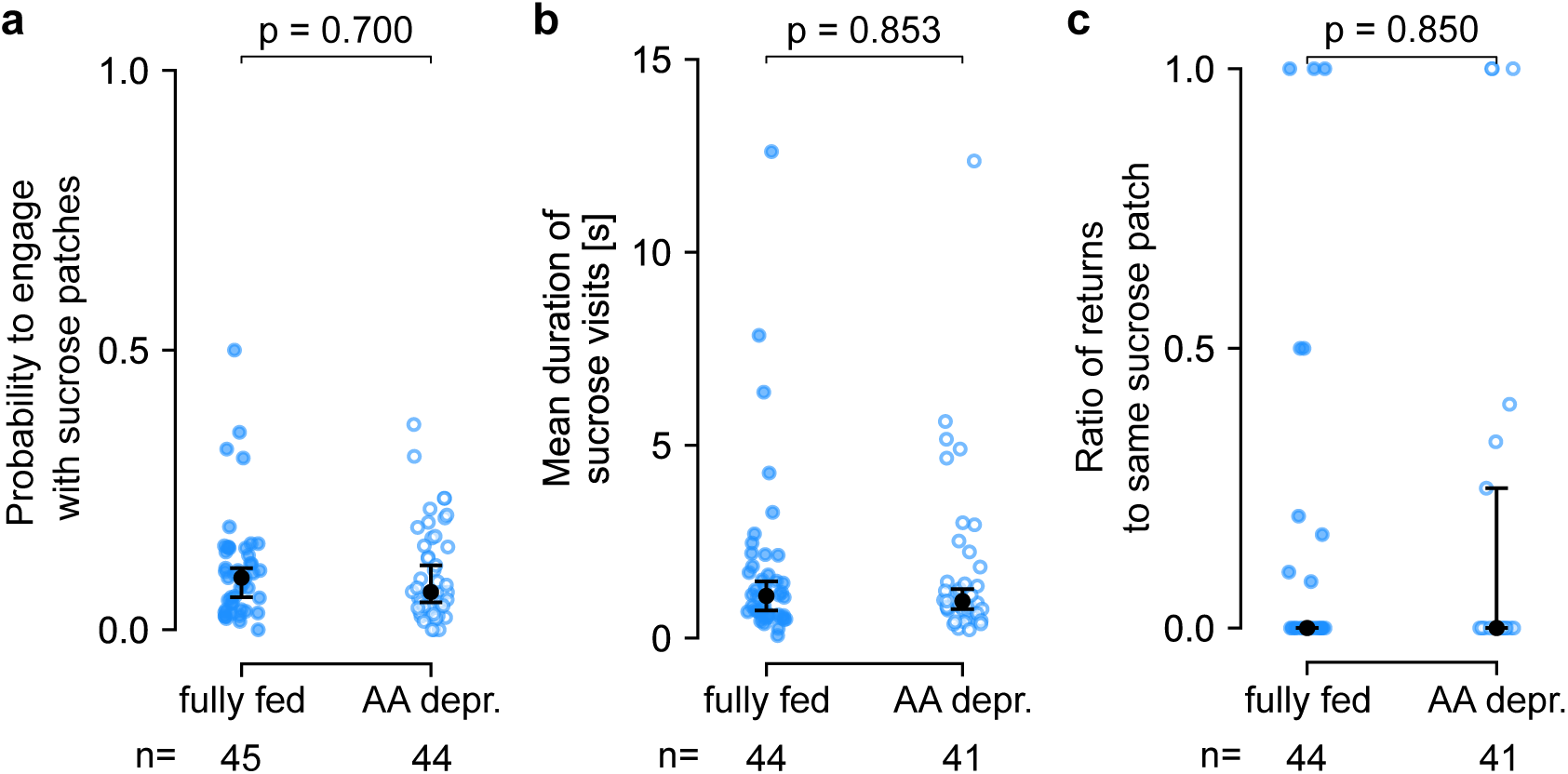
**Amino acid (AA) deprivation does not modulate foraging decision parameters for sucrose patches. a**, Probability to engage with sucrose patches for fully fed and AA deprived flies. **b**, Mean duration of sucrose visits for fully fed and AA deprived flies. **c**, Ratio of returns to the same sucrose patch for fully fed and AA deprived flies. **a-c**, Data points are individual flies, black dots and bars indicate median, and extrema within 1.5 of the inter-quartile range (IQR), respectively. P-values are obtained by performing Wilcoxon rank-sum tests, *n =* 41-45.

**Supplementary Figure 4.**
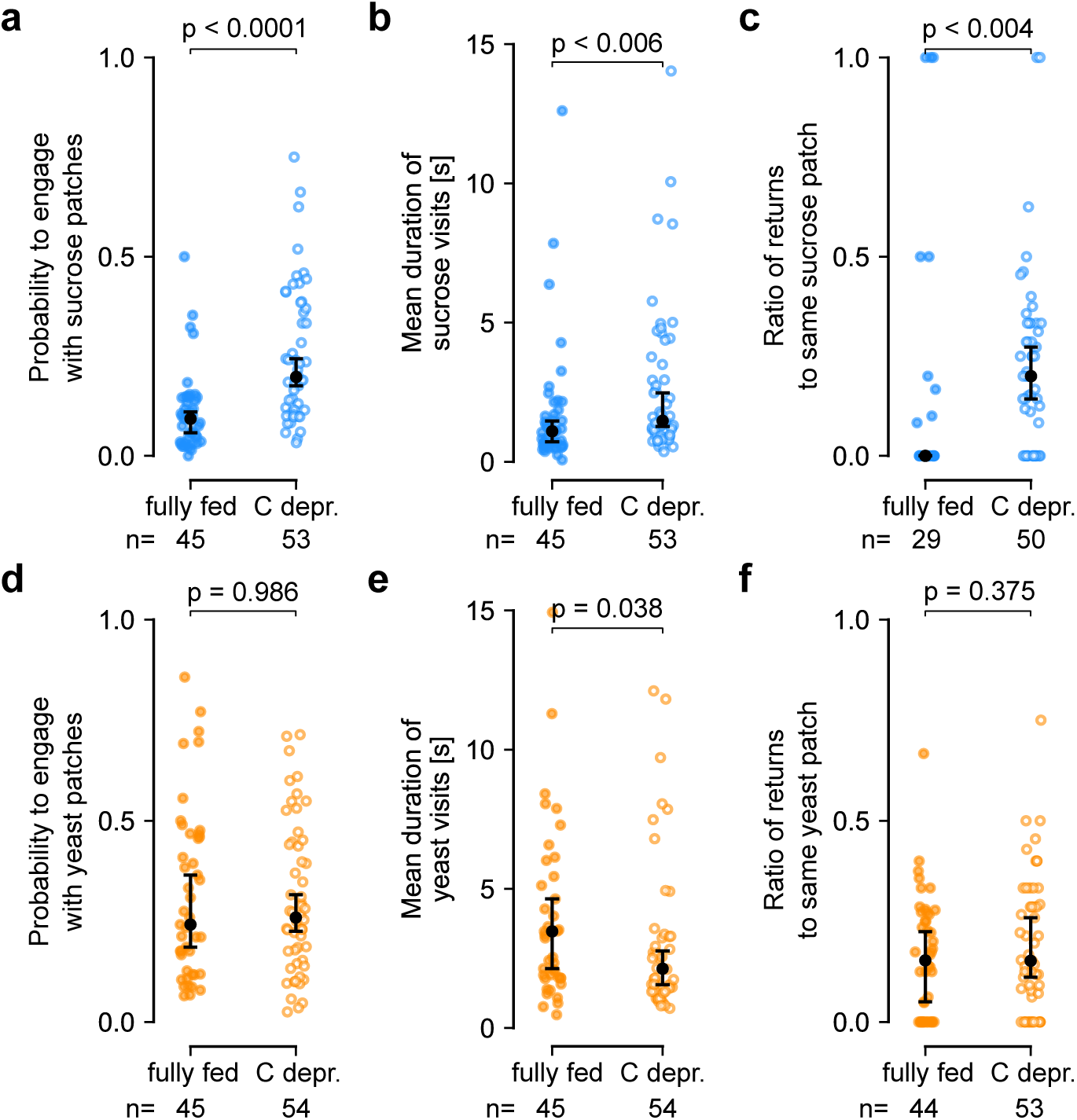
**Carbohydrate deprived (C depr.) flies adapt their foraging decision parameters on sucrose but not on yeast patches. a**, Probability to engage with sucrose patches for fully fed and carbohydrate deprived flies. **b**, Mean duration of sucrose visits comparing fully fed and carbohydrate deprived flies. **c**, Ratio of returns to the same sucrose patch for fully fed and carbohydrate deprived flies. **d**, Probability to engage with yeast patches for fully fed and carbohydrate deprived flies. **e**, Mean duration of yeast visits for fully fed and carbohydrate deprived flies. **f**, Ratio of returns to the same yeast patch for fully fed and AA deprived flies. **a-f**, Data points are individual flies, black dots and bars indicate median, and extrema within 1.5 of the inter-quartile range (IQR), respectively. P-values are obtained by performing Wilcoxon rank-sum tests, *n =* 29-45.

**Supplementary Figure 5.**
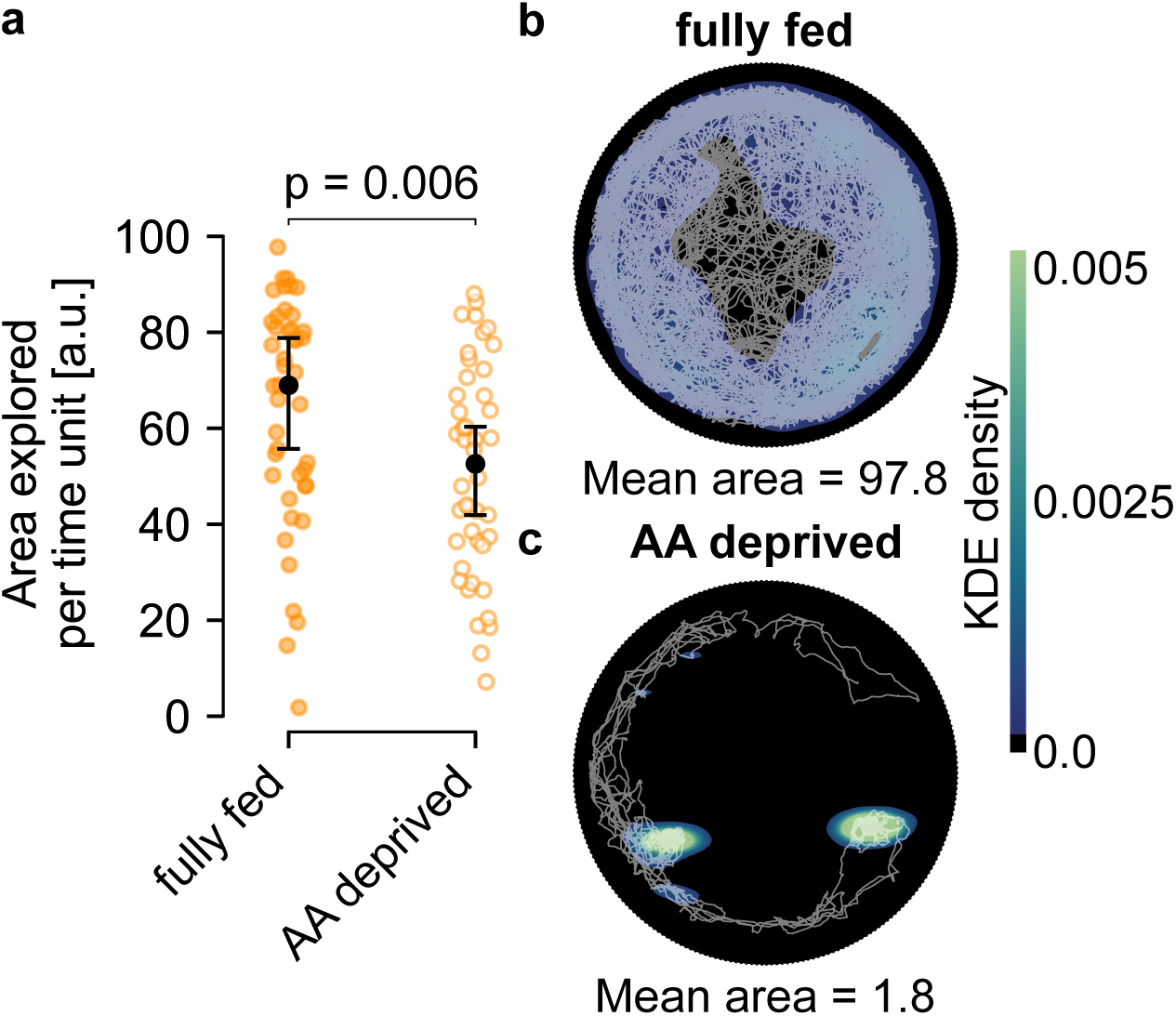
**AA deprived flies explore less area per time unit compared to fully fed flies. a**, Area explored per time unit for fully fed and AA deprived flies. Data points are individual flies, black dots and bars indicate median, and extrema within 1.5 of the inter-quartile range (IQR), respectively. P-values are obtained by performing Wilcoxon rank-sum tests, *n =* 44-45. **b-c**, Example trajectories of exploratory patterns for a fully fed **(b)** and an AA deprived **(c)** fly. Colormap indicates trajectory density based on Kernel Density Estimation (KDE). White lines are the trajectories of a representative, individual fly during the whole assay (1 hour).

**Supplementary Figure 6.**
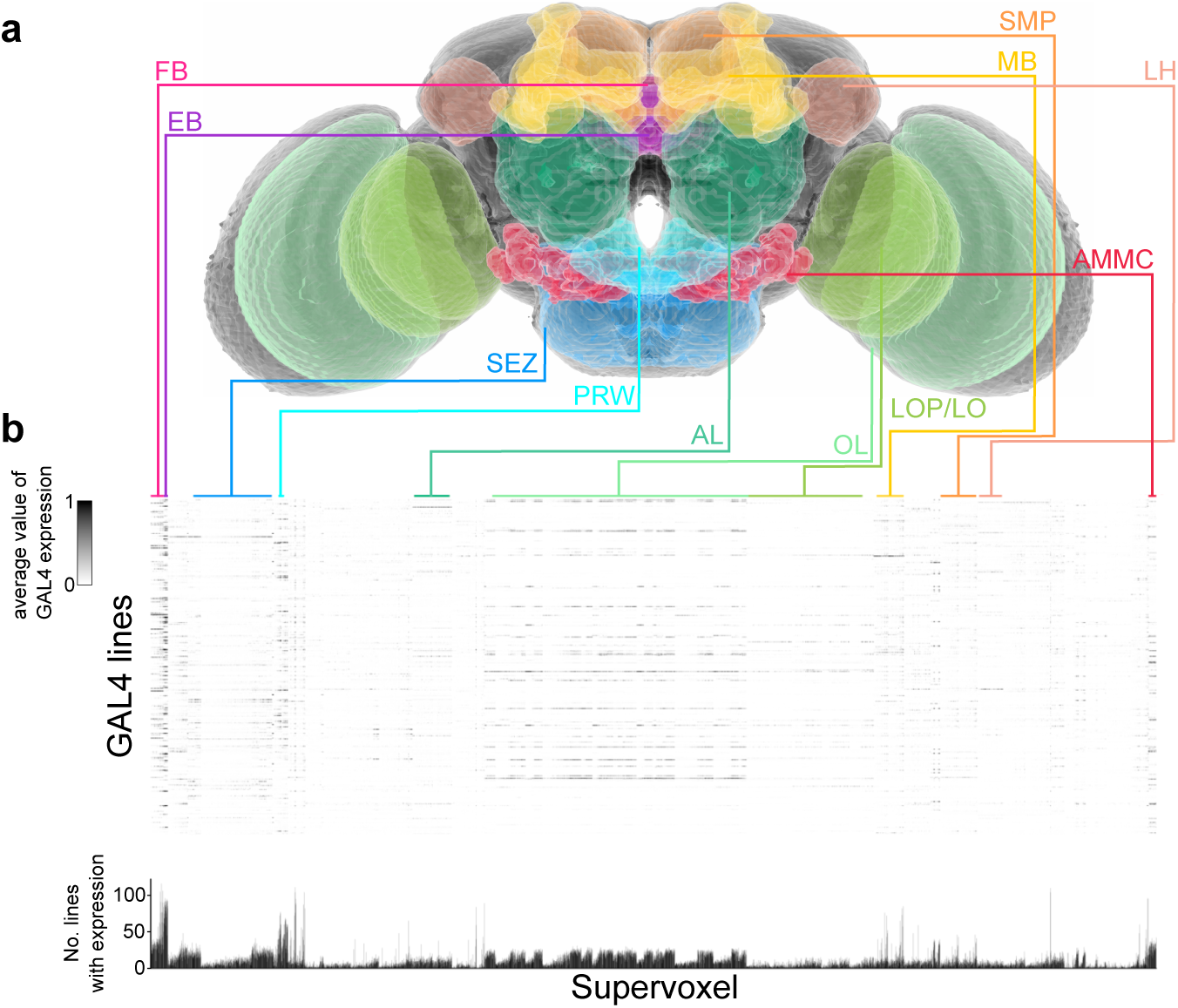
**GAL4 driver lines used in the large-scale silencing screen label diverse brain regions. a**, Image of neuropils targeted for the silencing screen rendered using Virtual Fly Brain (Court et al., 2023). FB = Fan-Shaped Body, EB = Ellipsoid Body, SEZ = Subesophageal Zone, PRW = Prow, AL = Antennal Lobes, OL = Optic Lobes, LOP/LO = Lobular Complex (Lobula Plate and Lobula), MB = Mushroom Bodies, SMP = Superior Medial Protocerebrum, AMMC = Antennal Mechanosensory and Motor Center, LH = Lateral Horns. **b**, Heatmap of supervoxels labeled by GAL4 driver lines used in the silencing screen. Every row corresponds to one GAL4 line. Anatomical supervoxel definitions were taken from Robie et al. (2017). Bottom panel is a histogram of the total number of lines with expression in the corresponding supervoxels.

**Supplementary Figure 7.**
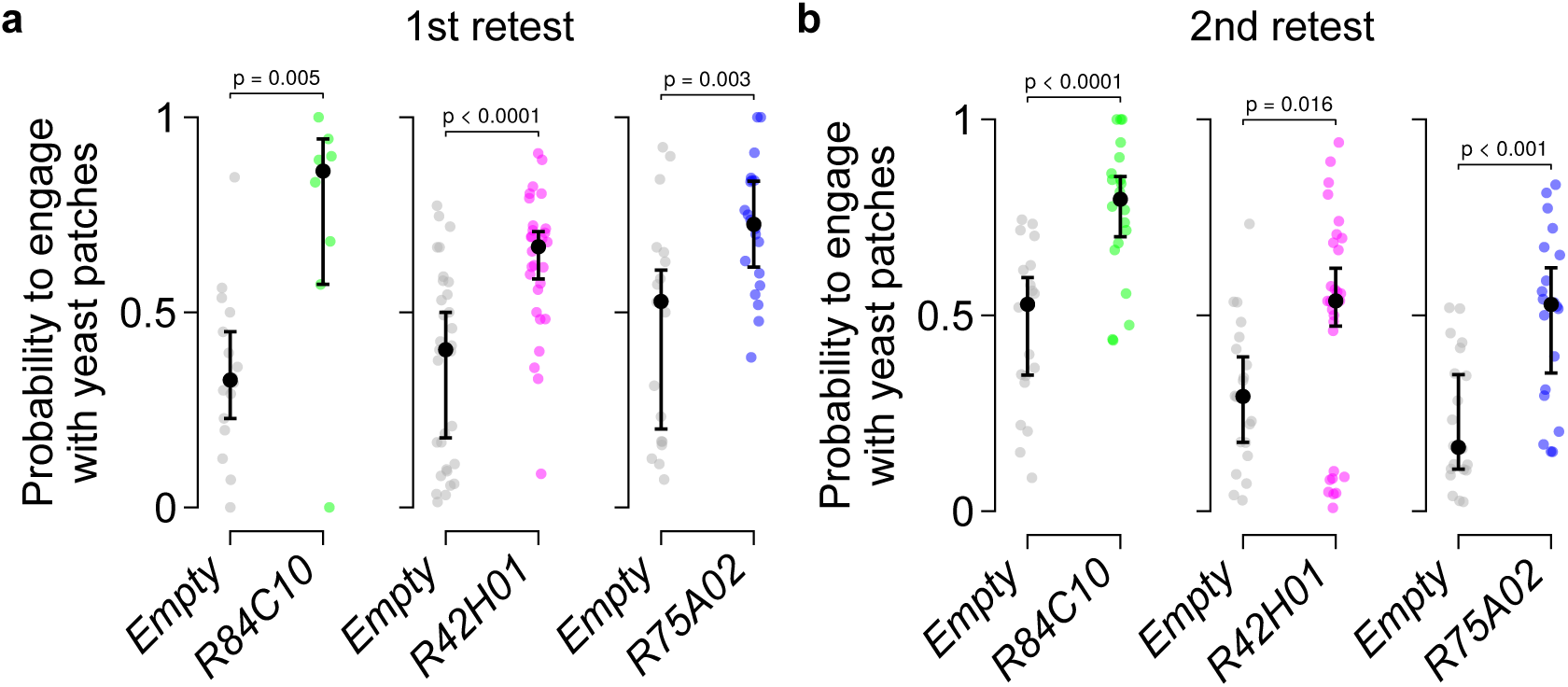
**Silencing GAL4 lines labeling dFB neurons shows a reproducible increase in the probability of engaging with a yeast patch. a-b**, Effect of silencing neurons labeled by *R84C10-*, *R42H01-*, and *R75A02-GAL4* on the probability to engage with yeast patches in the **(a)**, first and **(b)** second retest. Data points are individual flies, black dots and bars indicate median, and extrema within 1.5 of the inter-quartile range (IQR), respectively. P-values are obtained by performing Wilcoxon rank-sum tests, *n =* 8-20.

**Supplementary Figure 8.**
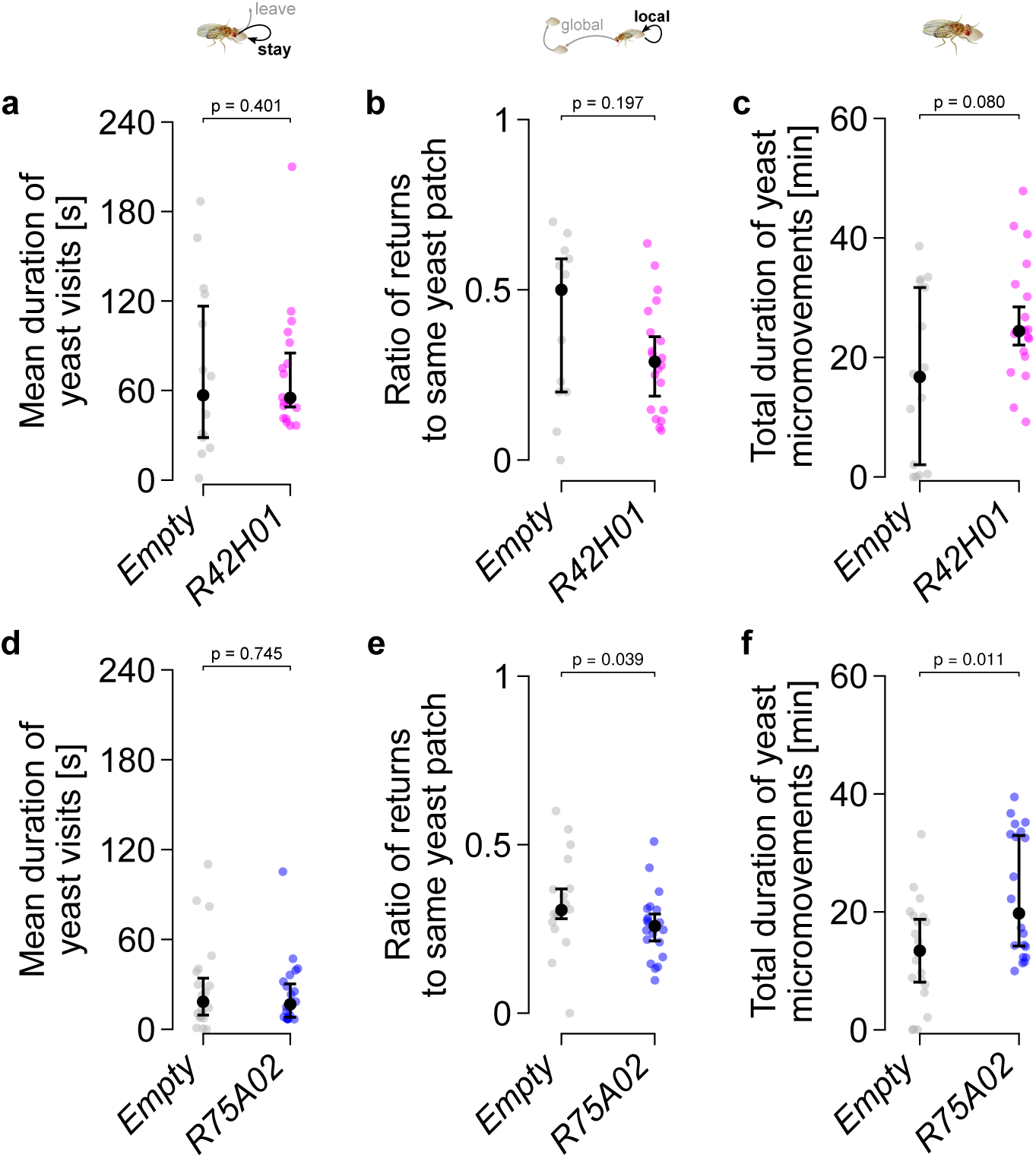
Silencing GAL4 lines labeling dFB neurons does not strongly modulate other foraging parameters. Effect of silencing neurons labeled by **a**, *42H01-*, and **d**, *R75A02-GAL4* on the mean duration of yeast visits. Effect of silencing neurons labeled by **b**, *R42H01-*, and **e**, *R75A02-GAL4* on the ratio of returns to the same yeast patch. Effect of silencing neurons labeled by **c**, *R42H01-*, and **f**, *R75A02-GAL4* on the total duration of yeast micromovements. **a-f**, Data points are individual flies, black dots and bars indicate median, and extrema within 1.5 of the inter-quartile range (IQR), respectively. P-values are obtained by performing Wilcoxon rank-sum tests, *n =* 17-20.

**Supplementary Figure 9.**
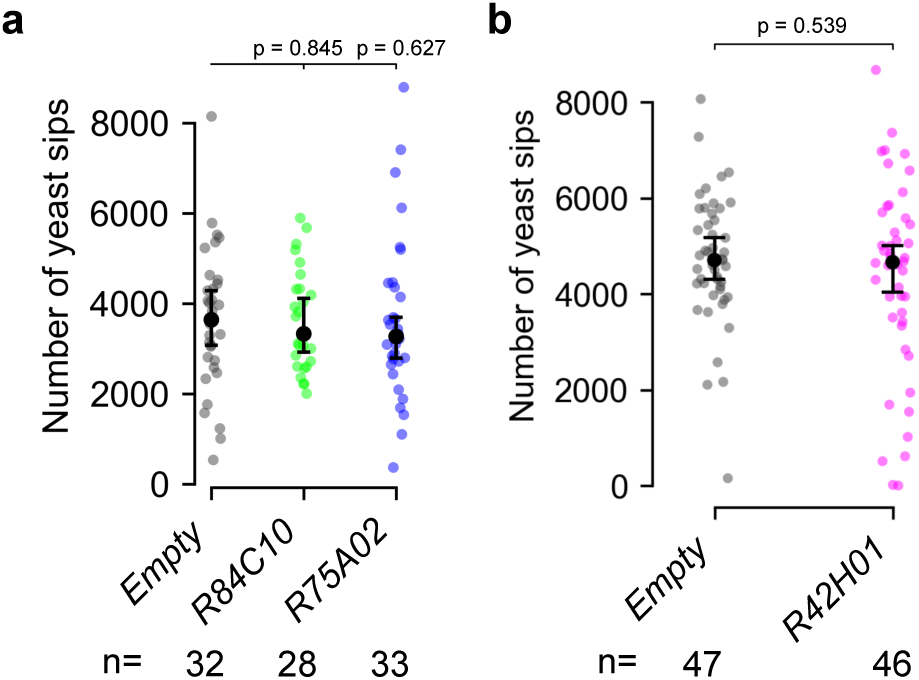
**Yeast feeding as measured using the flyPAD is not changed upon silencing dFB neurons. a**, Effect of silencing neurons labeled by *R84C10-* and *R75A02-GAL4* on the number of yeast sips on yeast as measured using the flyPAD setup. **b**, Effect of silencing neurons labeled by *R42H01-GAL4* on the number of yeast sips as measured using the flyPAD setup. **a-b**, Data points are individual flies, black dots and bars indicate median, and extrema within 1.5 of the inter-quartile range (IQR), respectively. P-values are obtained by performing Wilcoxon rank-sum tests, *n =* 28-47.

**Supplementary Figure 10.**
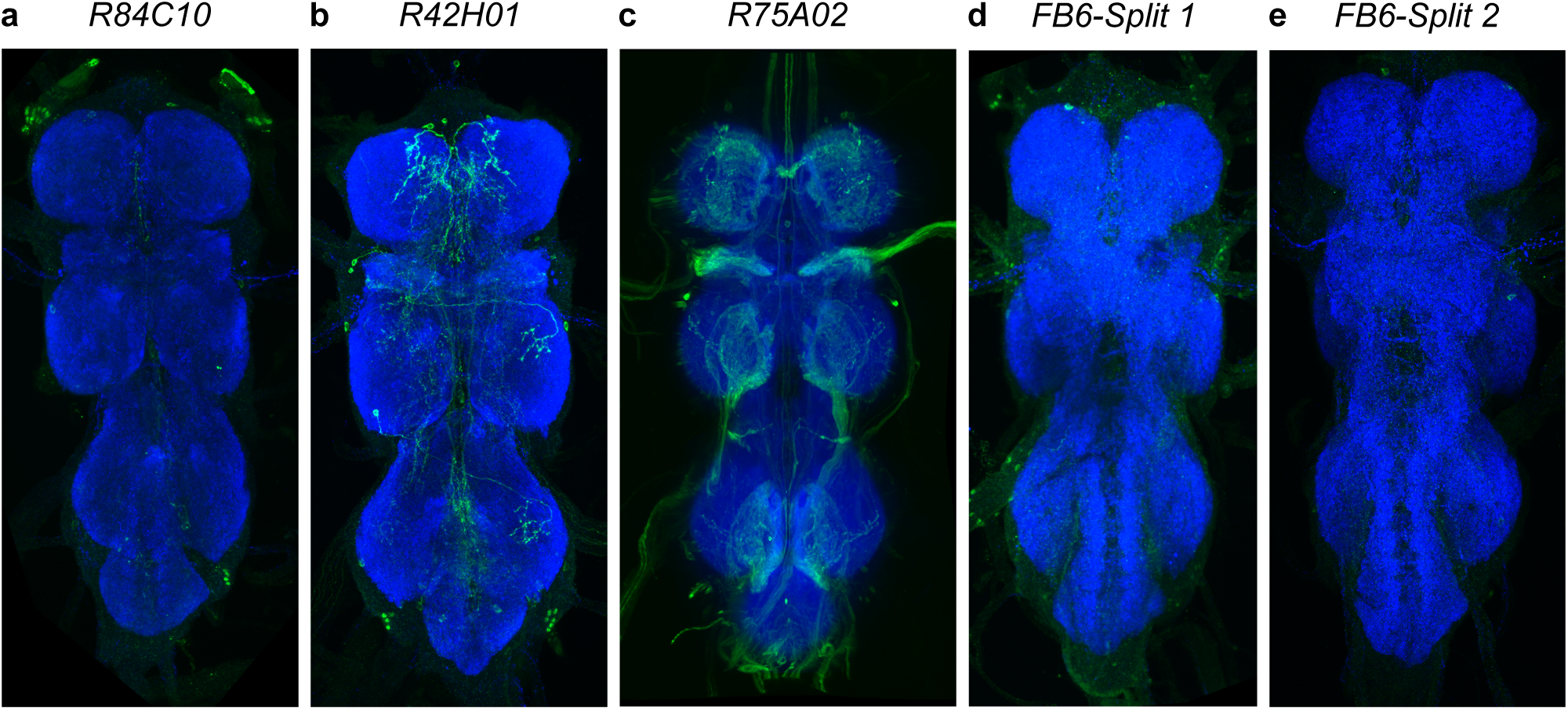
**Expression patterns of different GAL4 and Split-GAL4 driver lines in the ventral nerve cord. a-e**, Maximum intensity projections of confocal stacks for GFP expression in the Ventral Nerve Cords (VNCs) driven by **(a)** *R84C10-*, **(b)** *R42H01-*, **(c)** *75A02-*, **(d)** *FB6 Split 1-*, and **(e)** *FB6 Split 2-GAL4*.

**Supplementary Figure 11.**
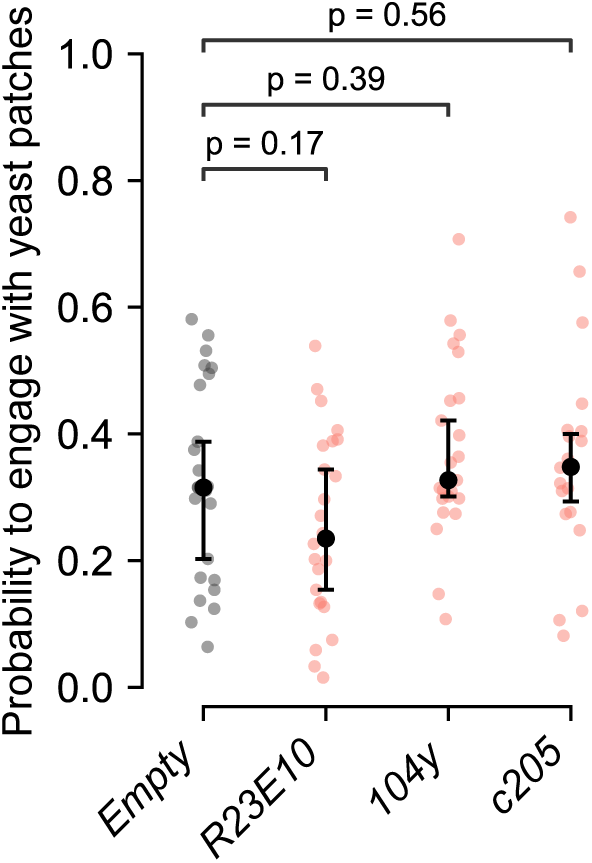
Silencing sleep-inducing GAL4 driver lines labeling dFB neurons does not affect the probability to engage with yeast patches. Effect of silencing neurons labeled by *R23E10-*, *104y-*, and *c205-GAL4* on the probability to engage with yeast patches. Data points are individual flies, black dots and bars indicate median, and extrema within 1.5 of the inter-quartile range (IQR), respectively. P-values are obtained by performing Wilcoxon rank-sum tests, *n =* 20-24.

**Supplementary Figure 12.**
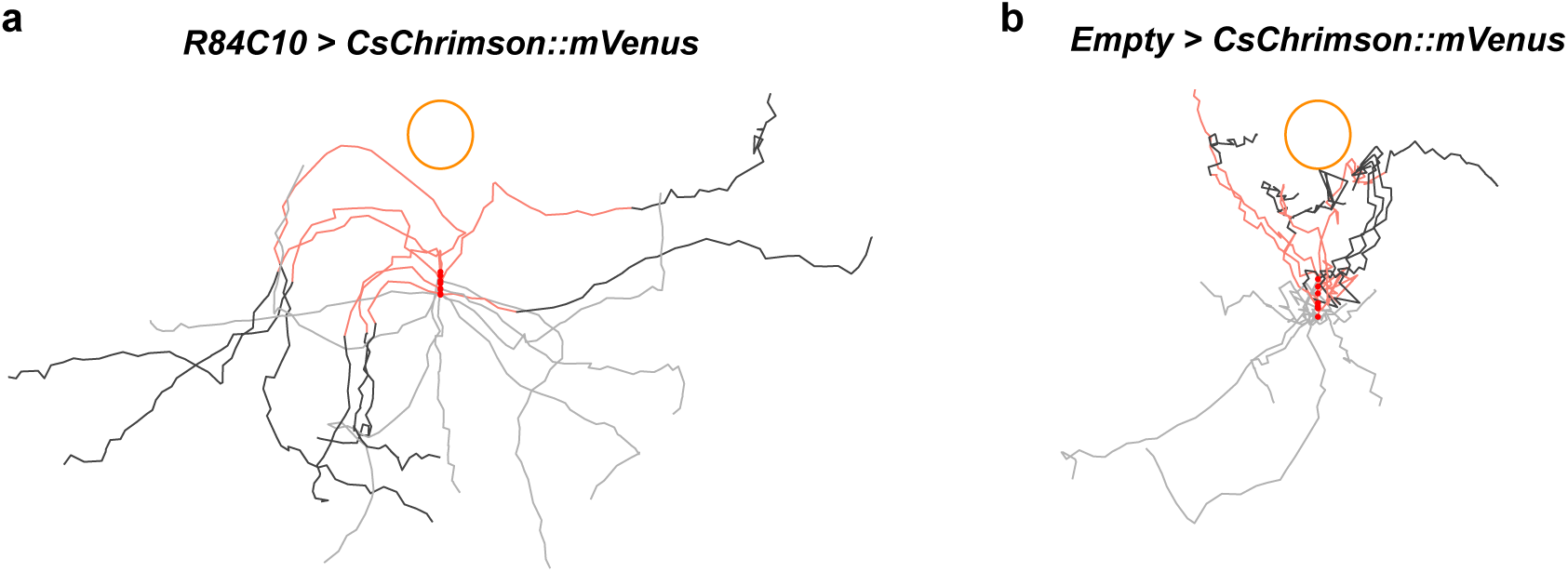
Optogenetic activation of dFB neurons during food patch approach leads flies to turn away from food patch. Sample trajectories of flies expressing *CsChrimson::mVenus* under the control of the *R84C10-GAL4* driver **(a)** or control flies **(b)**. LED light was triggered when the fly entered the area within a 1 cm radius around the yeast patch. The orange circle indicates the position and edge of the yeast patch. Trajectories were rotated with respect to the yeast patch such that the locations of the stimulation onset are overlapping. Light gray: trajectory before LED onset. Red: trajectory during the LED stimulation (1 second). Dark gray: trajectory after the end of LED stimulation. The onset of the LED activation for each trajectory is indicated with a red dot.

**Supplementary Figure 13.**
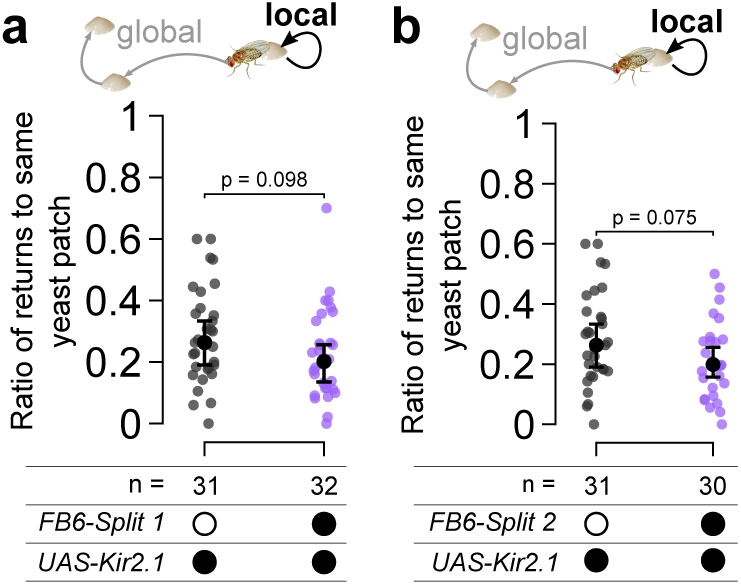
**Silencing FB6A neurons does not consistently modulate other foraging parameters. a-b**, Effect of silencing neurons labeled by *FB6-Split 1-* **(a)** and *FB6-Split 2-GAL4* **(b)** on the ratio of returns to the same yeast patch. Data points are individual flies, black dots and bars indicate median, and extrema within 1.5 of the inter-quartile range (IQR), respectively. P-values are obtained by performing Wilcoxon rank-sum tests, *n =* 30-32.

**Supplementary Figure 14.**
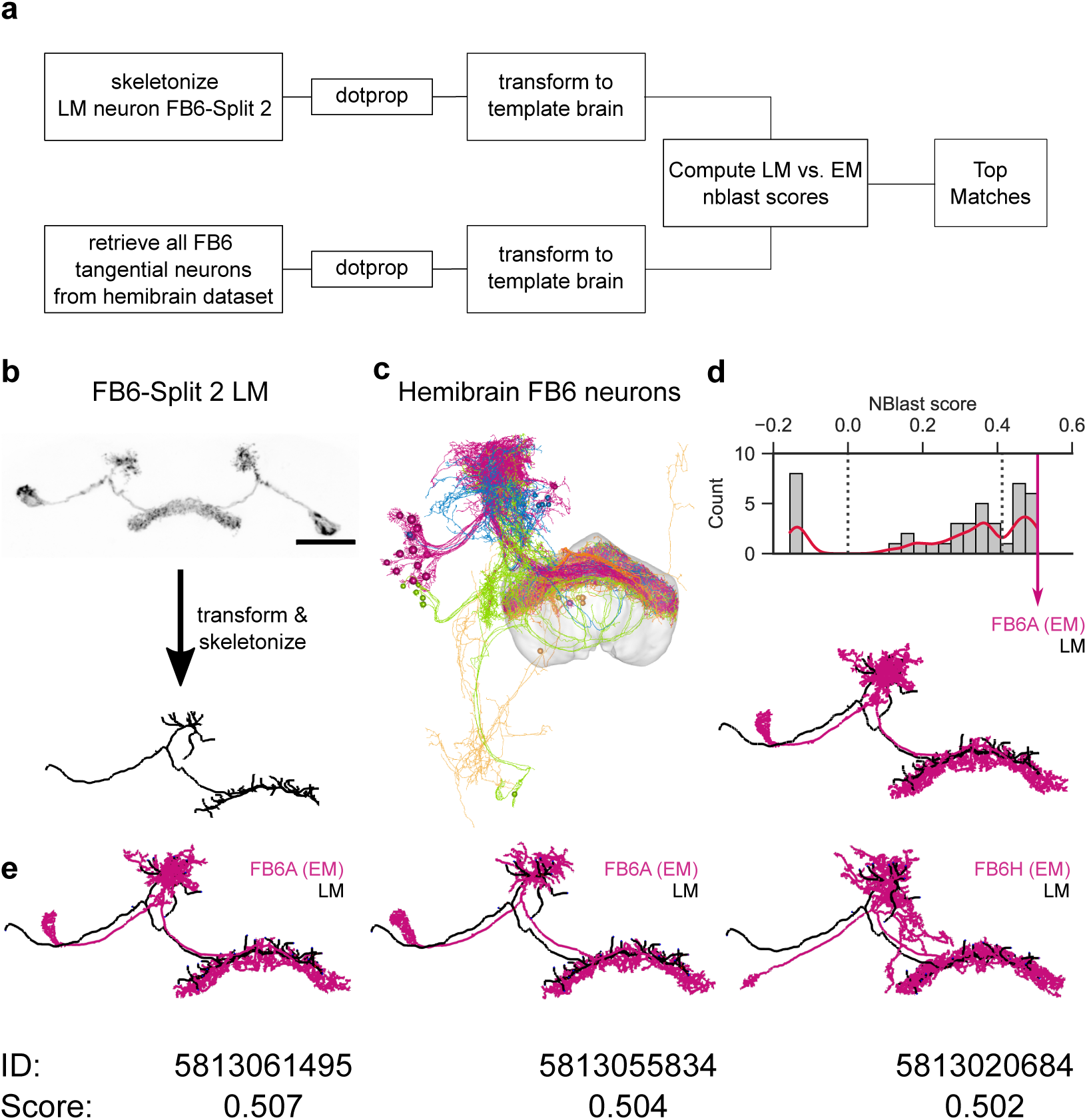
**Computational pipeline comparing morphological similarities across microscopy modalities reveals FB6-Split driver lines labeling FB6A. a**, The workflow used to compare LM and EM skeletons and identify top matches for the neurons labeled in the light microscopy dataset with neurons in the EM dataset. **b**, The confocal image obtained using the *FB6-Split 2-GAL4* driver line was registered to a template brain (JFRC2), skeletonized, and converted to dotprop representation. **c**, Rendering of all FB6 tangential neurons identified in the hemibrain dataset with the FB neuropil rendered in a transparent gray mesh. **d**, Histogram for the similarity scores obtained by comparing the LM skeleton to each of the hemibrain FB6 tangential neurons using NBLAST (Costa et al., 2016). The FB6A neuron (the EM and LM skeletons are rendered below) had the highest NBLAST score. **e**, Rendering of the EM skeletons (with the corresponding hemibrain skeleton ID and NBLAST scores below) of the three most similar neurons according to NBLAST scores (magenta) and the LM skeleton of the FB6 neuron (black).

**Supplementary Figure 15.**
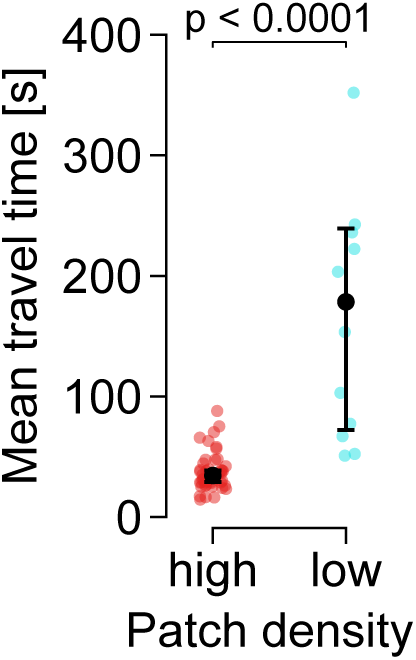
Mean travel time is longer in the low patch density arena. Mean travel time in seconds of wildtype flies foraging in high or low patch density arenas. Data points are individual flies, black dots and bars indicate median, and extrema within 1.5 of the inter-quartile range (IQR), respectively. P-values are obtained by performing Wilcoxon rank-sum tests, *n =* 12-44.

**Supplementary Figure 16.**
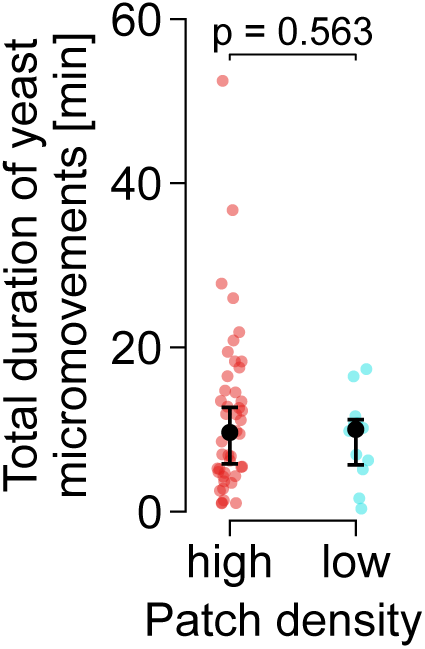
Total duration of yeast micromovements is not modulated by patch density. Total duration of yeast micromovements in minutes for wildtype flies foraging in high or low patch density arenas. Data points are individual flies, black dots and bars indicate median, and extrema within 1.5 of the inter-quartile range (IQR), respectively. P-values are obtained by performing Wilcoxon rank-sum tests, *n =* 12-44.

**Supplementary Figure 17.**
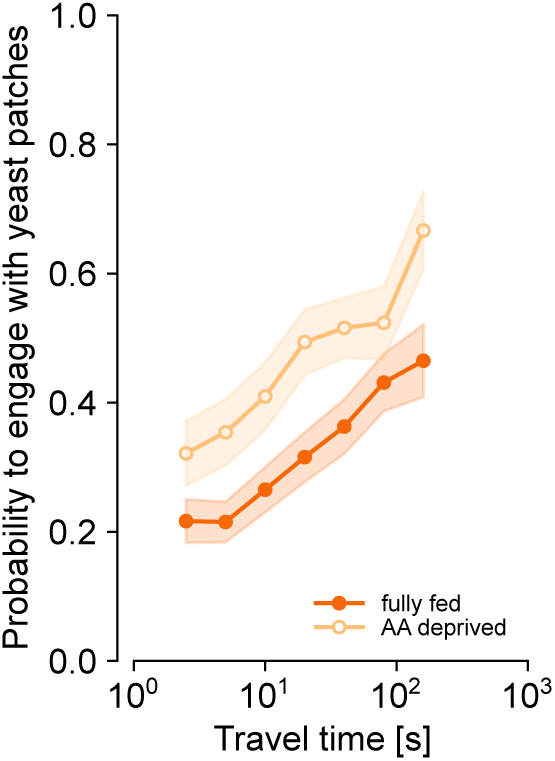
Protein state shifts the correlation between patch engagement and travel time towards a higher engagement probability but does not abolish the slope of the relationship. Probability to engage with yeast patches for different bins of travel times prior to the patch encounter for fully fed and AA-deprived flies, respectively. Data points and lines indicate mean across the flies and shaded areas indicate mean*±*sem, *n =* 44-45.

**Supplementary Video 1. Protein state changes the foraging behavior of flies.** Video of a fully fed (left) and an AA-deprived fly (right) foraging for 6 minutes (18x speed) in an arena containing yeast and sucrose food patches. The color of the trajectories change towards warmer colors over time.

**Supplementary Video 2. Activating dFB neurons specifically reduces the probability of the fly to transition from exploration to exploitation.** Example trajectories showing the effect of individual LED stimulations (red dots) upon approaching the food patch for experimental (*R84C10>CsChrimson*) and control flies (*Empty*).

